# A conserved RNA switch for acetylcholine receptor clustering at neuromuscular junctions in chordates

**DOI:** 10.1101/2024.07.05.602308

**Authors:** Md. Faruk Hossain, Sydney Popsuj, Burcu Vitrinel, Nicole A. Kaplan, Alberto Stolfi, Lionel Christiaen, Matteo Ruggiu

## Abstract

In mammals, neuromuscular synapses rely on clustering of acetylcholine receptors (AChRs) in the muscle plasma membrane, ensuring optimal stimulation by motor neuron-released acetylcholine neurotransmitter. This clustering depends on a complex pathway based on alternative splicing of *Agrin* mRNAs by the RNA-binding proteins Nova1/2. Neuron-specific expression of Nova1/2 ensures the inclusion of small “Z” exons in *Agrin,* resulting in a neural-specific form of this extracellular proteoglycan carrying a short peptide motif that is required for binding to Lrp4 receptors on the muscle side, which in turn stimulate AChR clustering. Here we show that this intricate pathway is remarkably conserved in *Ciona robusta,* a non-vertebrate chordate in the tunicate subphylum. We use *in vivo* tissue-specific CRISPR/Cas9-mediated mutagenesis and heterologous “mini-gene” alternative splicing assays in cultured mammalian cells to show that *Ciona* Nova is also necessary and sufficient for *Agrin* Z exon inclusion and downstream AChR clustering. We present evidence that, although the overall pathway is well conserved, there are some surprising differences in Nova structure-function between *Ciona* and mammals. We further show that, in *Ciona* motor neurons, the transcription factor Ebf is a key activator of *Nova* expression, thus ultimately linking this RNA switch to a conserved, motor neuron-specific transcriptional regulatory network.

## INTRODUCTION

The human brain contains about 86 billion neurons (Azevedo et al., 2009). For the brain to function, neurons have to communicate with each other via connections called synapses, and a single neuron can have tens of thousands of synapses (DeFelipe et al., 2002). This staggering architectural and circuitry complexity is essential to process sensory information and control the body’s response to external stimuli and, ultimately, cognition and behavior (Sanes and Lichtman, 1999; Südhof, 2008). At the same time, such complexity makes it difficult to study individual synapses, particularly at the molecular level. Due to its large size and experimental accessibility, the neuromuscular junction (NMJ), a peripheral cholinergic synapse between a motor neuron and a muscle cell (Hall and Sanes, 1993; Sanes and Lichtman, 1999, 2001; Slater, 2017) is arguably the best-understood mammalian synapse, and NMJ deterioration is at the center of the neuromuscular disorder amyotrophic lateral sclerosis (Cappello and Francolini, 2017; Dupuis and Loeffler, 2009; Fischer et al., 2004; Mitchell and Borasio, 2007; Pasinelli and Brown, 2006). Motor neurons secrete a large basal lamina proteoglycan named Agrin for its ability to promote aggregation of acetylcholine receptor (AChR) clusters on the muscle surface (Gautam et al., 1996; Nitkin et al., 1987; Reist et al., 1992). Agrin is synthesized by most cells of the body, but only neurons produce an alternatively spliced isoform of *Agrin* termed Z^+^ (or neural) *Agrin*. The two Z microexons encode a short domain of 8-19 amino acids that confers up to a 1,000-fold increase in AChR clustering activity compared to Z^-^ Agrin, the isoform that does not include the Z exons (Bezakova et al., 2001; Burgess et al., 1999; Ferns et al., 1993; Gautam et al., 1996; Gesemann et al., 1996; Gesemann et al., 1995; Hoch et al., 1993). In fact, *Agrin* KO mice and mice in which the Z exons have been deleted both die at birth from diaphragmatic paralysis (Burgess et al., 1999; Gautam et al., 1996), suggesting that the Z exons are essential for Agrin function. Interaction of Z^+^ Agrin with the Agrin postsynaptic receptor LDLR-related protein 4 (Lrp4)(Kim et al., 2008; Weatherbee et al., 2006; Zhang et al., 2008) leads to the phosphorylation of the muscle-specific receptor tyrosine kinase MuSK, and, through a cascade of events, it induces AChR clustering on the muscle (DeChiara et al., 1996; Gautam et al., 1996; Glass et al., 1996; Glass and Yancopoulos, 1997; Ruegg and Bixby, 1998), and defects in this signaling pathway are responsible for the congenital neuromuscular disorder Congenital Myasthenic Syndrome, or CMS (Beeson et al., 2006; Beeson et al., 2008; Ben Ammar et al., 2013; Bogdanik and Burgess, 2011; Chevessier et al., 2004; Engel et al., 2008; Engel and Sine, 2005; Hamuro et al., 2008; Huzé et al., 2009; Maselli et al., 2010; Maselli et al., 2012; Müller et al., 2004; Müller et al., 2006; Nicole et al., 2014; Ohkawara et al., 2014; Ohkawara et al., 2020; Rudell et al., 2019; Wang et al., 2020).

Despite the central role of *Agrin* Z exons in synapse biology, only recently have we started to understand how Z exon splicing is regulated. The neuron-enriched splicing factors NOVA1 and NOVA2 underlie an autoimmune neuromuscular disorder (Buckanovich et al., 1993; Darnell and Posner, 2003), and *NOVA2* mutations cause a severe form of neurodevelopmental disorder (Mattioli et al., 2020). Double knockout mice for both *Nova1* and *Nova2* fail to include the Z exons of *Agrin* (Ruggiu et al., 2009); however, as Nova proteins regulate about 700 alternative splicing events in the brain (Zhang et al., 2010), whether this is a direct effect. How Nova proteins bind to and promote *Agrin* Z exon inclusion is still largely unknown.

To elucidate the role of Nova in regulation of *Agrin* Z exon splicing, we focused on the tunicate *Ciona robusta*. Tunicates, or sea squirts, are the closest living relatives to vertebrates within the chordate phylum (Delsuc et al., 2006; Putnam et al., 2008). The central nervous system (CNS) of the *Ciona* larva contains only 177 neurons (Ryan et al., 2016), and its connectome has recently been completed (Ryan et al., 2016, 2017, 2018; Ryan and Meinertzhagen, 2019). Yet this minimal nervous system is formed and compartmentalized in a very similar manner as the larger nervous systems of vertebrates (Hudson, 2016). This relative cellular simplicity, alongside rapid development and a compact genome that has not undergone duplications seen in vertebrates, makes *Ciona* uniquely suited to dissect the evolutionary biology of protein-RNA regulatory switches that are important for synapse biology and neurologic disorders. In this work we show that the motor neuron terminal selector Ebf (Kratsios et al., 2012) activates the transcription of *Nova*, which is present as a single copy gene in *Ciona*. Nova protein in the larval motor neurons directly promotes the inclusion of *Agrin* Z exons, which in turn stimulates acetylcholine clustering at the NMJ through Lrp4 receptors just as in vertebrates. By elucidating this splicing event at the molecular level, we uncover unexpected features of Nova that contribute to its splicing regulation function. We also provide evidence of coevolution of Nova and the regulatory sequences embedded in the *Agrin* pre-mRNA that mediate Nova-dependent splicing, revealing “developmental system drift” of an otherwise highly conserved RNA splicing-dependent molecular switch.

## RESULTS

### Identification of divergent Z exons in *Ciona robusta Agrin*

Previous bioinformatic analysis of potential Nova splicing targets in *Ciona* and other invertebrates did not indicate *Agrin* as a potential target, suggesting that *Agrin* Z exon splicing regulation by Nova was a vertebrate-specific innovation (Hrus et al., 2007; Irimia et al., 2011). However, we identified two cryptic exons in between annotated exons 40 and 41 (**Figure 1A,B**), which were confirmed by cloning from mixed embryonic stage cDNA library. These were named “Z6” and “Z5” as they were found to encode 6- and 5 amino acid-long polypeptide sequences, respectively (**Figure 1C**).

**Figure 1.**
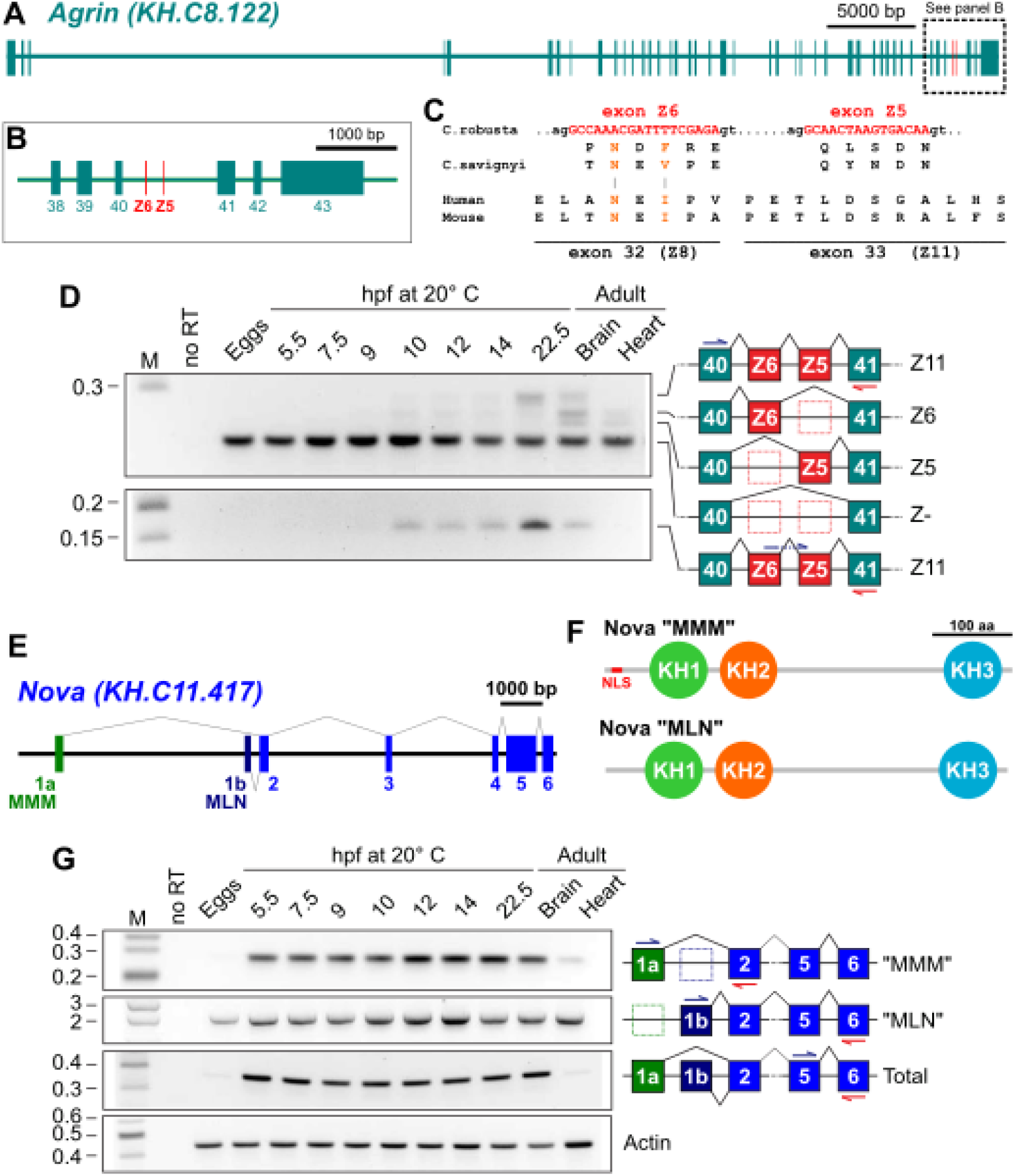
*Agrin* and *Nova* expression and splicing in *Ciona robusta* development. A) Diagram of *Agrin* gene (KyotoHoya gene model ID: KH.C8.122) in *C. robusta* showing exons as thicker rectangles. B) Zoomed in view of dashed region spanning constitutive exons 38-43 of *Agrin*, indicating the position of the Z exons (Z6 and Z5). C) Predicted DNA coding sequences (top) of *C. robusta* Z6 and Z5 exons of *Agrin,* showing predicted protein sequences underneath, aligned to corresponding *Ciona savignyi* protein sequences. At bottom, protein sequences encoded by the Z8 and Z11 exons of mouse and human *Agrin,* for comparison. Proposed conserved NxI/V/F peptide motif highlighted in orange font. D) RT-PCR gel profiling Z exon inclusion in *C. robusta Agrin* mRNAs extracted from different developmental stages and two different adult tissues. Top gel performed with primers specific to flanking constitutive exons 40 and 41, which can simultaneously amplify Z-negative and different Z+ isoforms (Z5, Z6, Z11). Bottom gel performed using a forward primer spanning the Z5/Z6 exon-exon junction, which amplifies only the Z11 isoform. E) Diagram of *Nova* gene (gene ID KH.C11.417) in *C. robusta,* indicating the two alternative 1^st^ exons (1a and 1b) encoding the Nova proteins starting with the peptides MMM and MLN, respectively. F) Diagrams of the predicted protein domain organization “MMM” and “MLN” isoforms of Nova, showing the N-terminal NLS present only in the MMM isoform. G) RT-PCR profiling of Nova expression and alternative splicing, using primers amplifying either MMM, MLN, or all Nova isoforms. Cellular actin transcripts used as a positive control for RT-PCR. M: DNA molecular weight marker. no RT: no reverse transcriptase added. Embryonic developmental stages given in hours post-fertilization (hpf) at 20°C.

Exon Z6 in particular was found to encode an N-X-F motif that might be functionally equivalent to the N-X-I/V motif that is encoded by the Z8 exon (exon 32) of mammals and mediates the interaction between neural Agrin and Lrp4 (Guarino et al., 2019; Zong et al., 2012). Through predicted protein sequence alignments, we found that the corresponding motif in the related species *C. savignyi* is N-X-V, supporting the idea that these sequences are likely to be conserved, functional motifs for Lrp4 binding encoded by homologous Z exons.

To determine when these Z exons are included in the *Agrin* mRNAs during development, we performed a time-series of RT-PCR using primers specifically designed to amplify the region encoded by the Z6 or Z5 exons (**Figure 1D**). Although *Agrin* transcripts were detected in unfertilized eggs and early embryonic stages, Z exon-specific amplicons were only detected starting around 10 hours post-fertilization (hpf) at 20°C (∼stage 22, or mid-tailbud II), continuing through larval stages. “Z11” *Agrin* transcripts (containing both Z6 and Z5 exons) were detected from 10 hpf onwards, including in the adult brain but not in the heart (**Figure 1D**). These data suggest that *Agrin* Z exon inclusion is occurring primarily in neural tissue, and during neuronal differentiation in embryogenesis.

### Developmental regulation of *Nova* and *Agrin* expression in the Ciona embryo

To determine whether *Nova* might be expressed at the same time when we observe Z exon inclusion in *Agrin* transcripts, we performed a similar RT-PCR time-series for the single ortholog of mammalian *Nova1/Nova2* in *C. robusta.* This gene, which we call simply “*Nova”,* appears to encode two major isoforms that differ in their first exon (**Figure 1E**). Transcripts including the more 5’ first exon (exon “1a”) encode an isoform of the Nova protein that includes a predicted N-terminal nuclear localization signal (NLS). In contrast, those including the more 3’ first exon (exon “1b”) do not appear to encode an NLS. We termed these two isoforms “MMM” and “MLN” (**Figure 1F**), respectively, based on the first three amino acid residues of their protein sequences. By RT-PCR we detected both isoforms as early as the unfertilized eggs, though the “MLN” isoform appeared to be the most abundant one at this stage. Both transcript variants were expressed throughout embryogenesis and in the adult stage, though expression appeared more abundant in the brain than in the heart. In larvae and adult brains, both isoforms appeared to be equally abundant (**Figure S1**).

*Nova* expression during *Ciona* development was previously investigated using whole-mount mRNA *in situ* hybridization (ISH) and reported as specific to the CNS starting at the neurula stage onwards (Irimia et al., 2011). However, the exact identities of Nova- expressing cells were not reported. Therefore, we decided to characterize *Nova* expression in greater detail. By ISH, we first detected *Nova* transcription in neural progenitors at the early gastrula stage (**Figure 2A**). *Nova* transcription continued in neural progenitors in the neural plate at late gastrula (**Figure 2B**), and subsequently throughout the neural tube in early mid-tailbud stage embryos (**Figure 2C**). At this stage we also noticed expression in non-neural tissues: cardiopharyngeal mesoderm (e.g. trunk ventral cells, or TVCs), a subset of mesenchyme cells, and very weakly in oral siphon muscle precursors and posterior endoderm (**Figure 2C**). The cardiopharygeal mesoderm staining confirms earlier reports of *Nova* expression in this lineage by microarray and single-cell RNAseq profiling (Christiaen et al., 2008; Razy-Krajka et al., 2014; Vitrinel et al., 2023; Wang et al., 2019).

**Figure 2.**
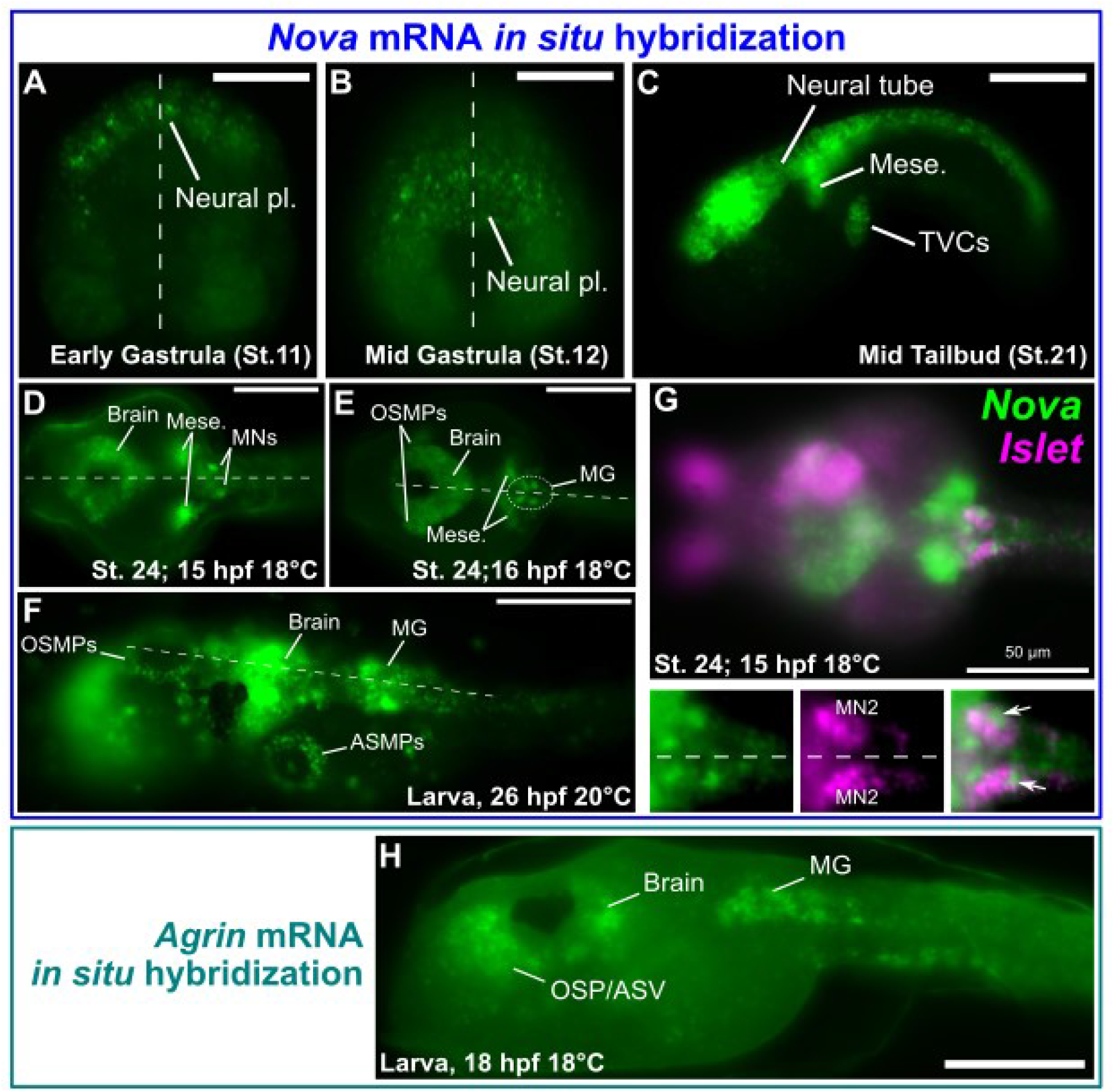
Expression of *Nova* and *Agrin* detemined by *in situ* hybridization. A) Whole-mount fluorescent mRNA *in situ* hybridization (ISH) of *Nova* in *C. robusta,* showing earliest detectable zygotic expression in neural plate progenitors at the early gastrula stage (Hotta Stage 11, ∼4.5 hours post-fertilization (hpf) at 20°C. B) Expression is observed in the nascent neural plate at Stage 12 (∼5 hpf at 20°C). C) *Nova* is expressed throughout the neural tube and also in mesoderm-derived mesenchyme and trunk ventral cells (TVCs, also known as cardiopharyngeal progenitors) at mid-tailbud stage, Stage 21 (∼9.5 hpf at 20°C). D) Expression of *Nova* is now observed to be maintained/upregulated in motor neurons of the Motor Ganglion (MG), as well as in the brain/posterior sensory vesicle and mesenchyme in earlier Stage 24 embryos (∼15 hpf at 18°C). E) *Nova* is also seen to be activated in oral siphon muscle progenitors (OSMPs) in later Stage 24 embryos (∼16 hpf at 18°C). G) Two-color double ISH of *Nova* (green) and *Islet* (magenta) showing co-expression in the bilaterally symmetric Motor Neuron 2 (MN2) pair of cells (arrows). Note that *Nova* mRNA distribution appears more nuclear than *Islet* in these cells at this stage. H) ISH of *Agrin* in the *C. robusta* larva showing expression in the oral siphon/anterior sensory vesicle (OSP/ASV) region, in the larval brain, and in the Motor Ganglion (MG). Dashed lines in any panel indicate embryonic midline in dorsal views. All scale bars = 50 µm.

In later tailbud embryos (∼stage 24), we detected *de novo* upregulation of *Nova* transcripts in specific left/right pairs of cells in the motor ganglion (MG), which appeared to be differentiating motor neurons (**Figure 2D,E**). First, at 15 hpf at 18°C, *Nova* was upregulated in a pair of posterior cells (**Figure 2D**). Slightly later (16 hpf, 18°C), *Nova* transcripts were observed in at least two pairs of MG cells (**Figure 2E**). In these cells, upregulation of *Nova* was observed as a strong pulse of stained transcripts localized primarily to the cell nucleus. *Nova* expression continued in the MG, brain, and siphon muscle precursors during the larval stage (**Figure 2F**).

To precisely identify the *Nova-*expressing cells in the MG, we performed double ISH for *Nova* and *Islet* (Giuliano et al., 1998; Imai et al., 2009), a known marker of the “Motor Neuron 2” pair of motor neurons (MN2) that form *en passant* synapses at sites of AChR clusters in the tail muscles (Nishino et al., 2011). Double ISH for *Nova* and *Islet* revealed co-expression in MN2 at ∼stage 24 (15 hpf at 18°C, **Figure 2G**). The identification of these posterior-most *Nova*+ cells as motor neurons was confirmed by performing ISH for *Nova* in embryos electroporated with *Fgf8/17/18>H2B::mCherry* plasmid (**Figure S2**), which marks the A9.30 lineage of Ciona (Imai et al., 2009). It has been shown that MN2 cells are derived from the A9.32 lineage and are invariantly positioned immediately posterior to the A9.30 lineage (Navarrete and Levine, 2016; Stolfi and Levine, 2011). Indeed, we detected *Nova* expression in the cell just posterior to the A9.30 lineage, not co-expressed with H2B::mCherry, confirming its expression in MN2 (**Figure S2**). Finally, ISH also revealed that *Agrin* is transcribed throughout the MG, in addition to other cells around the larval brain and sensory vesicle (**Figure H**).

Taken together, these data show that *Nova* and *Agrin* are co-expressed in larval neurons, in particular the motor neurons that form NMJs with the muscles of the tail.

### A minigene assay to study regulation of *Ciona Agrin* splicing in cell culture

Given their co-expression in *Ciona* neurons, we investigated whether Nova might also promote inclusion of the Z exons during alternative splicing of *Agrin* in *Ciona,* as Nova1/2 proteins do in vertebrates (Ruggiu et al., 2009). To do this, we developed a *Ciona Agrin* minigene splicing assay based on similar assays previously described (Gaildrat et al., 2010; Smith and Lynch, 2014; Stoss et al., 1999). Plasmids encoding exons 40, 41, and the intervening introns and Z exons under the *cis*-regulatory control of the CMV promoter were co-transfected with different concentrations of *Ciona* or mouse Nova expression plasmids into cultured mammalian cells (**Figure 3A**). The inclusion of the Z exons in the resulting *Ciona Agrin* mini-transcripts was then assayed by RT-PCR on cDNA prepared from transfected cells. As expected, inclusion of *Ciona Agrin* Z exons increased linearly with increased dose of *Ciona* Nova (**Figure 3B**).

**Figure 3.**
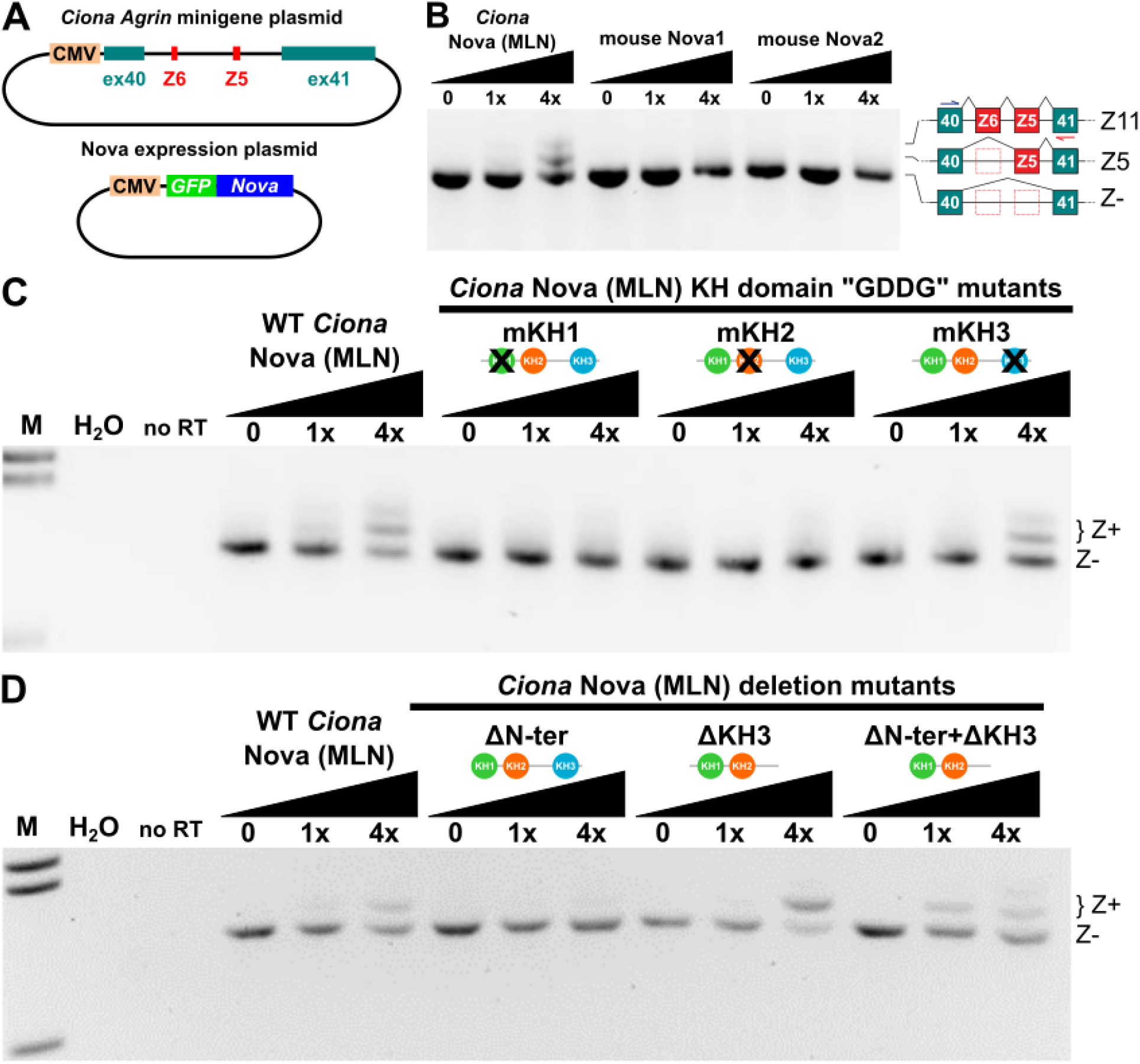
*Ciona Agrin* minigene Z exon inclusion assay. A) Diagram of *Ciona robusta Agrin* minigene plasmid (top) used for alternative splicing assays in cultured mammalian (HEK293T) cells, along with Nova expression plasmids (bottom). B) *Ciona* Nova (“MLN” isoform tested) can promote *Ciona Agrin* minigene Z exon inclusion, assayed by RT-PCR, while mouse Nova1 or Nova2 cannot. Identity of Z11, Z5, and Z- (Z-negative) confirmed by cloning and sequencing RT-PCR products. Z6 isoforms were not detected in the minigene assay. Black slope indicates increasing Nova expression plasmid dose. C) Testing the effect of “GDDG” mutations in each RNA-binding KH domain of *Ciona* Nova (KH1-3) using the same minigene assay as above. Abolishing the RNA-binding activity of KH1 and KH2, but not KH3, disrupts the ability of *Ciona* Nova to promote Z exon inclusion. D) Deleting the N-terminus of *Ciona* Nova (MLN isoform) abolishes its ability to promote *Agrin* Z exon inclusion in the minigene assay. This effect is nullified by concomitant deletion of the KH3 domain (see text for details). M: DNA molecular weight marker. H2O: using water instead of cDNA template for PCR. no RT: no reverse transcriptase added.

Curiously, only Z11 and Z5 isoforms were detected (**Figure 3B**), confirmed by cloning and sequencing, suggesting inclusion of the Z6 exon alone might be regulated by additional factors or sequences not present in our minigene assay. Unexpectedely, mouse Nova1 and Nova2 were unable to promote Z exon inclusion in the *Ciona Agrin* mini-transcripts (**Figure 3B**). This suggests that, although the regulation of *Agrin* splicing by Nova proteins might be conserved from tunicates to vertebrates, there may have been additional co-evolution that has resulted in divergent *cis/trans* compatibility: only *Ciona* Nova, not vertebrate Nova1/2, might be capable of splicing *Ciona Agrin*.

In vertebrates, Nova1/2 have the ability to bind pre-mRNAs through their three KH domains, which are all conserved in *Ciona* Nova (**Figure 1F**). However, it is not currently known which KH domains in Nova might mediate *Agrin* Z exon inclusion. Different KH domains of Nova1/2 can bind different RNA targets, resulting in complex mechanisms of binding and splicing by these proteins (Buckanovich and Darnell, 1997; Jensen et al., 2000; Teplova et al., 2011; Ule et al., 2006; Zhang et al., 2010). To test which KH domains of *Ciona* Nova are required for its ability to splice *Ciona Agrin* to include the Z exons, we used our minigene assay to test different KH domain mutants of the more ubiquitous “MLN” isoform of Nova. The three KH domains of *Ciona* Nova were disrupted (individually or in combination) by changing the G-X-X-G loop sequence to G-D-D-G, which impairs RNA binding without affecting domain stability (Hollingworth et al., 2012). According to our assay, we determined that the KH1 and KH2 domains of *Ciona* Nova are required for optimal Z exon inclusion, while disrupting the KH3 domain did not appear to have any noticeable effect (**Figure 3C**, **Figure S3**). Surprisingly, deleting the short N-terminus of *Ciona* Nova alone also abolished its ability to promote Z exon inclusion (**Figure 3D**). This effect was rescued by deleting the KH3 domain, even though the KH3 deletion on its own did not affect Z exon inclusion (**Figure 3D**). Based on these data, we propose that the N-terminus of *Ciona* Nova is a regulatory domain that inhibits the KH3 domain, allowing the protein to switch from a KH3- to a KH1/KH2- based splicing mode.

Finally, we asked whether there were any *cis*-regulatory sequences in the *Agrin* pre-mRNAs that might be important for its Nova-dependent splicing and Z exon inclusion. Indeed, we identified the presence of 18 potential Nova binding sites (YCAY) in the intron between exons Z5 and 41 (**Figure 4A**), with no other YCAY sequences present elsewhere in this region. As vertebrate Nova proteins have been shown to bind pre-mRNAs via intronic YCAY clusters (Dredge and Darnell, 2003; Dredge et al., 2005; Jelen et al., 2007; Jensen et al., 2000), we tested whether disrupting these sequences in the *Ciona Agrin* minigene plasmids might block the ability of *Ciona* Nova to promote Z exon inclusion. Indeed, we found that generating point mutations in some of these YCAY clusters greatly suppressed the inclusion of *Ciona Agrin* Z exons by *Ciona* Nova (**Figure 4B**, **Figure S4**). A footprint analysis indicated that the most crucial clusters mapped to YCAY sites 3-7 and 11-13 in the intron between the Z5 exon and constitutive exon 41 (**Figure 4C**). Therefore it appears *Ciona* Nova uses two Nova Intronic Splicing Enhancers, or NISEs (Dredge and Darnell, 2003), to promote Z exon inclusion, which requires at least two YCAY sequences in each element. Taken together, our minigene data suggest that *Ciona* Nova is capable of promoting the alternate splicing of *Agrin* pre-mRNAs through direct interactions between its KH1/KH2 domains and the intronic YCAY clusters in its target.

**Figure 4.**
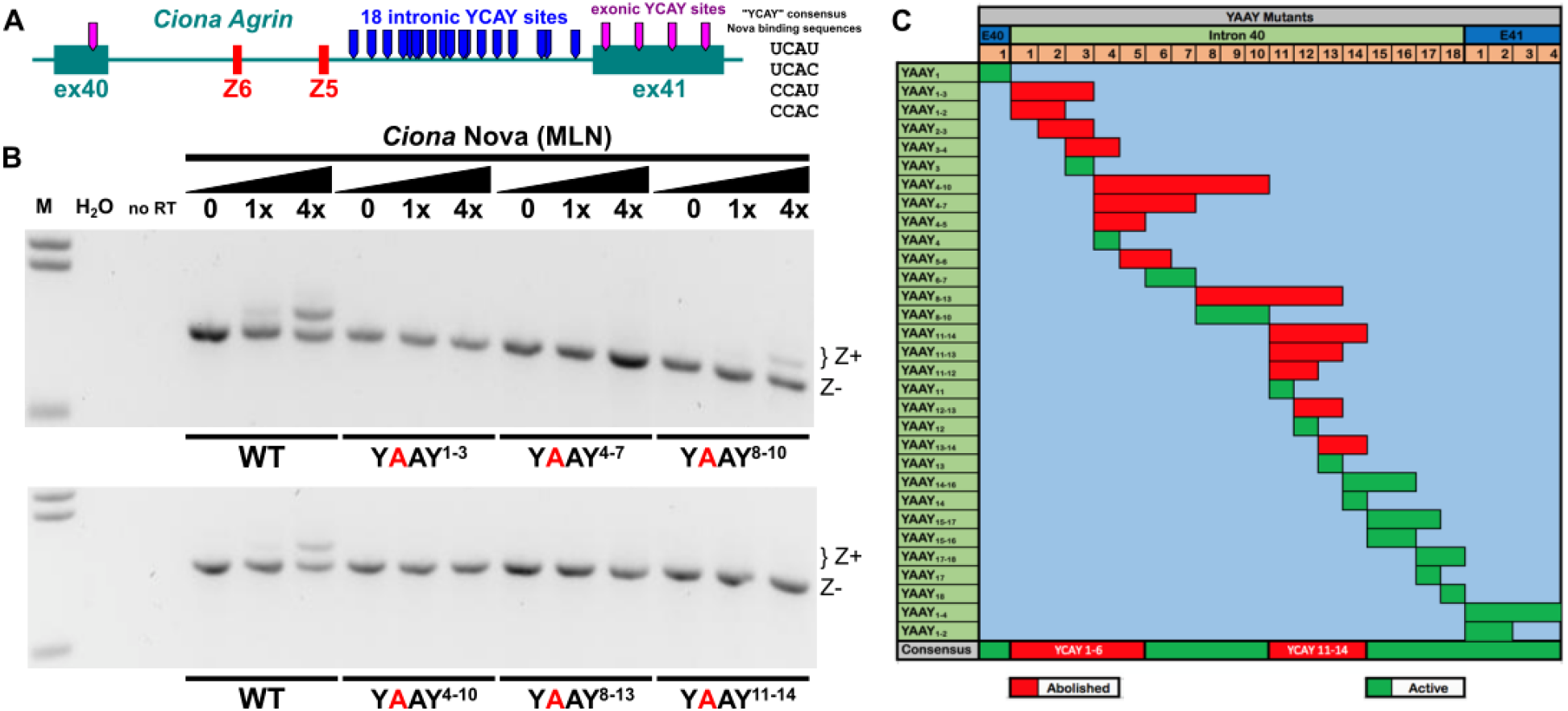
Minigene assay to test *cis*-regulatory elements that promote Z exon inclusion. A) Diagram of the Z exon region of *Ciona robusta Agrin,* indicating 18 potential Nova binding sites (consensus: YCAY) in the intron between exon Z5 and constitutive exon 41, as well as exonic YCAY sequences (magenta tabs). B) *Ciona Agrin* minigene Z exon inclusion detected by RT-PCR, using minigenes bearing different combination of candidate Nova-binding site mutations (YCAY>YAAY) predicted to disrupt Nova binding. Full set of mutations assayed shown in **Figure S4**. M: molecular weight marker. H2O: using water in place of cDNA template for PCR. no RT: no reverse transcriptase added. C) Chart summarizing effect of disrupting different YCAY sites in the *Ciona Agrin* minigene. Two clusters of intronic YCAY sites appear to be required for proper Nova-dependent Z exon inclusion: YCAY sites 1-6 and 11-14. Other intronic sites and sites in constitutive exons 40 or 41 are not required for Z exon inclusion.

### A conserved Agrin-Lrp4 pathway for AChR clustering at the NMJ

In mammals, Z+ Agrin released by MNs expressing Nova promotes AChR clustering in target muscles by binding to Lrp4 (Kim et al., 2008; Ruggiu et al., 2009; Zhang et al., 2008). Thus, we sought to test the potentially conserved role of Nova in regulating *Agrin* alternative splicing and downstream neuromuscular synapse development in *Ciona.* To do this, we turned to tissue-specific CRISPR/Cas9 (Gandhi et al., 2018). To first establish the role of Z+ Agrin in controlling the clustering of AChRs in the larval tail muscles at *en passant* synapses formed by MN2 (Nishino et al., 2011). We designed eight different single-chain guide RNAs (sgRNAs) targeting sequences flanking the Z exons in *Ciona Agrin,* a region spanning exons 39-41. To test if CRISPR/Cas9 using these sgRNAs could suppress Z exon inclusion, we performed qPCR on cDNAs generated from embryos co-electroporated with *Eef1a>Cas9,* to drive ubiquitous Cas9 expression (Stolfi et al., 2014), together with different combinations of our *Agrin-* targeting sgRNA constructs. Three out of four such combinations resulted in reduced Z11+ *Agrin* amplification (**Figure 5B**). We selected the subset of sgRNAs that showed the highest and most specific reduction in Z11+ transcript levels (combination #1) for further investigation (**Figure 5A**).

**Figure 5.**
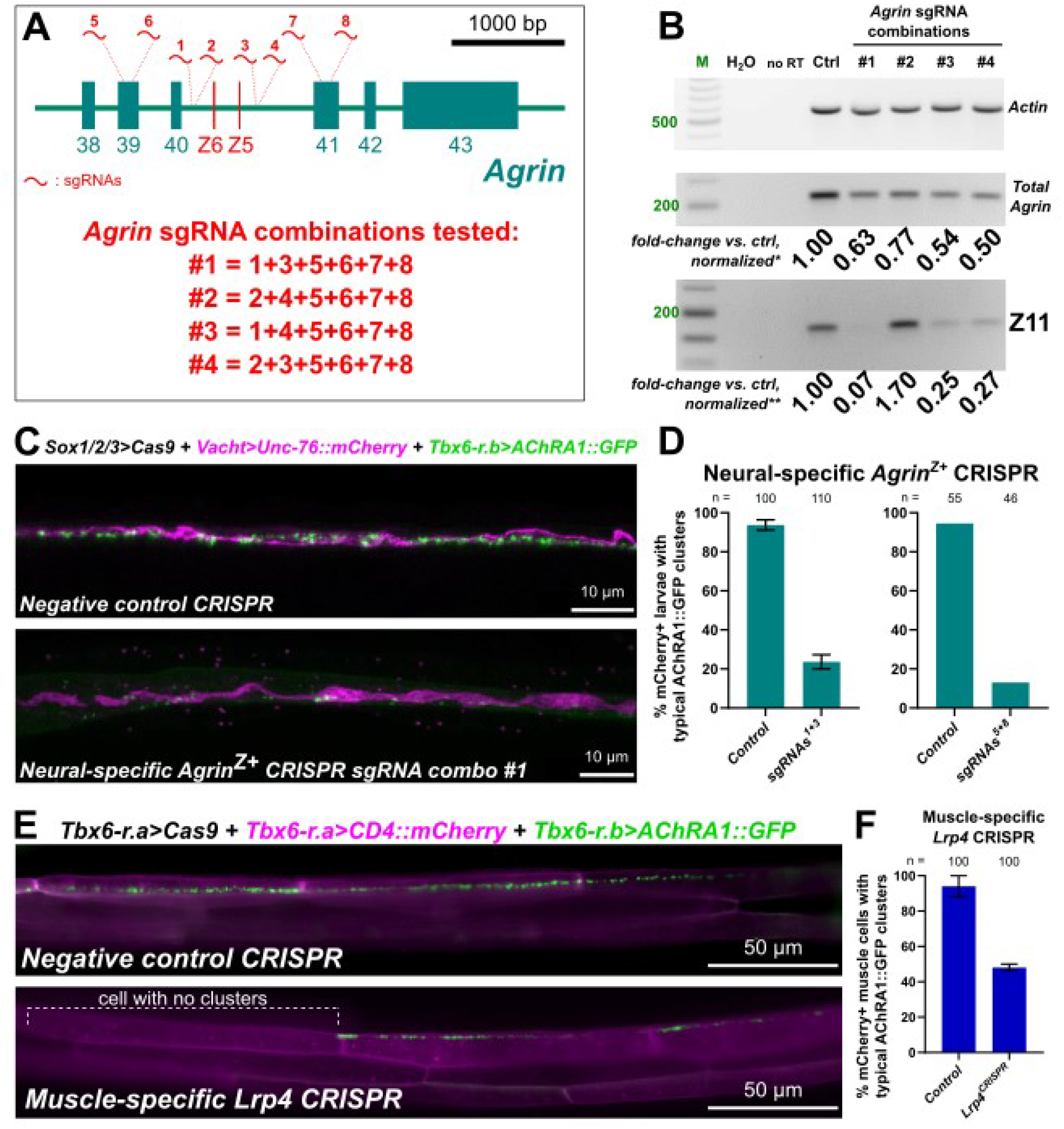
TIssue-specific CRISPR/Cas9-mediated mutagenesis of *Agrin* and *Lrp4.* A) Partial diagram of the *Agrin* gene from *Ciona robusta,* showing position of target sites and combinations of sgRNAs for CRISPR/Cas9-mediated mutagenesis. B) RT-PCR- based quantification of *Agrin* Z exon inclusion (from larvae collected at 22.5 hours post- fertilization at 20°C) following CRISPR-mediated disruption of sequences surrounding the Z exons. Selected sgRNA combinations are indicated in panel A, which were co- electroporated with *Eef1a>Cas9*. Fold-change of band intensity is compared to a negative control CRISPR sample. * total *Agrin* bands were normalized according to corresponding *Actin* band intensity. ** Z11 exon bands were normalized according to both *Actin* and total *Agrin* bands for each corresponding sample. M: molecular weight marker. H2O: water used in place of cDNA template for PCR. no RT: no reverse transcriptase added. C) Neural-specific CRISPR-mediated disruption of *Agrin* using sgRNAs indicated in the diagram above results in loss of Acetylcholine receptor A1::GFP clusters (AChRA1::GFP, green) in tail muscles, driven by the *Tbx6-r.b* promoter (Christiaen et al., 2009a). Motor neuron axons labeled by *VAChT>Unc- 76::mCherry* in magenta (Yoshida et al., 2004). D) Scoring loss of AChRA1::GFP clusters in the muscles upon targeting *Agrin* using more specific combinations of NGS- validated sgRNA pairs (1+3 and 5+8). 1+3 sgRNA pair tested in duplicate with at least 45 larvae in per condition and duplicate. Negative control larvae electroporated with negative control sgRNA instead. E) Muscle-specific CRISPR-mediated disruption (using *Tbx6-r.b>Cas9*) of the Agrin receptor-encoding gene *Lrp4* shows similar loss of AChRA1::GFP clusters. Effects were seen on a muscle cell-by-cell basis, as expected if the effect of disrupting the receptor is cell-autonomous. Negative control is actually muscle-specific CRISPR-mediated disruption of *Nova,* confirming neural-specific requirement of Nova as demonstrated further below in Figure 6. F) Scoring of larvae represented in the previous panel. Experiment performed in duplicate with 50 individual muscle cells examined per condition and duplicate.

We performed tissue-specific CRISPR/Cas9-mediated mutagenesis of the *Agrin* Z exon (*Agrin^Z+^*) region by co-electroporating the selected sgRNA combination #1 (sgRNAs 1, 3, 5, 6, 7, and 8) together with *Sox1/2/3>Cas9* plasmid. The *Sox1/2/3* promoter was used to drive Cas9 expression in neural progenitors, including the lineage that gives rise to the motor neurons of the *Ciona* larva (Stolfi et al., 2014). To assay AChR clustering at NMJs, we co-electroporated *Tbx6-r.b>AChRA1::GFP* plasmid to express in the tail muscles the AChRA1::GFP subunit fusion that was previously used to visualize such clusters postsynaptic to MN2 (Nishino et al., 2011). Neural-specific disruption of *Agrin^Z+^* significantly reduced AChRA1::GFP clusters in the tail muscles, at dorsal sites of contact with MN2 (**Figure 5C**, **Figure S5**). This effect was replicated using two different combinations of sgRNAs targeting more specifically the introns flanking the Z exons (sgRNAs 1 and 3), or exons 39 and 41 (sgRNAs 5 and 8)(**Figure 5D**), which were validated as generating indels at their appropriate target sites by genomic DNA amplicon sequencing (**Figure S6**).

To test whether Lrp4 might play a conserved role as a receptor for Agrin in the tail muscles of *Ciona,* we performed muscle-specific CRISPR/Cas9-mediated disruption. To target the *Lrp4* gene specifically in muscles, we co-electroporated *Tbx6-r.b>Cas9* together with validated *Lrp4*-targeting sgRNAs (**Figure S7**). Compared to the negative control, we found a significant reduction in the number of muscle cells with visible AChRA1::GFP clusters (**Figure 5E,F**). Often the AChRA1::GFP clusters were either present or entirely absent from whole muscle cells (**Figure 5E**), which in *Ciona* larvae are invariantly derived from different Tbx6-r.b+ precursors and therefore likely experience independent CRISPR/Cas9 mutagenesis events due to mosaic uptake of electroporated plasmids (Zeller et al., 2006). Taken together, these data suggest that Lrp4 is also required for AChR clustering in *Ciona* NMJs, similar to its role in vertebrates.

### *Ciona* Nova is required for *Agrin* Z exon inclusion and AChR clustering

We next asked whether in *Ciona* Nova plays a conserved role in splicing of neural-specific, Z+ *Agrin* mRNAs to induce AChR clustering at the NMJ. First, we designed and validated three sgRNAs targeting the *Nova* gene by CRISPR/Cas9 (**Figure 6A**, **Figure S8**). We then used RT-PCR to investigate *Agrin* Z exon inclusion upon CRISPR/Cas9- mediated disruption of *Nova* in *Ciona* larvae (**Figure 6B**). Z+ *Agrin* transcripts were significantly reduced upon co-electroporation of *Eef1a>Cas9* and any one of the three *Nova-*targeting sgRNAs individually, or in combination (**Figure S9**). This effect was reproduced in triplicate, while overexpression of Nova (*Eef1a>Nova*) resulted in increased Z exon inclusion (**Figure 6C**). Taken together, these results suggest that *Ciona* Nova is sufficient and necessary for *Agrin* Z exon inclusion *in vivo,* just like Nova1/2 in mammals (Ruggiu et al., 2009). When we looked at AChRA1::GFP clusters at NMJs, we saw that they were reduced in frequency or density in *Nova* CRISPR larvae compared to negative control larvae (**Figure 6D-F**). Furthermore, AChR clustering was partially rescued by expressing CRISPR-insensitive *Nova* cDNA in MN2 (**Figure 6E**). Taken together, these results reveal that a Nova-Agrin-Lrp4 pathway for AChR receptor clustering at the neuromuscular synapse is conserved from mammals to tunicates.

**Figure 6.**
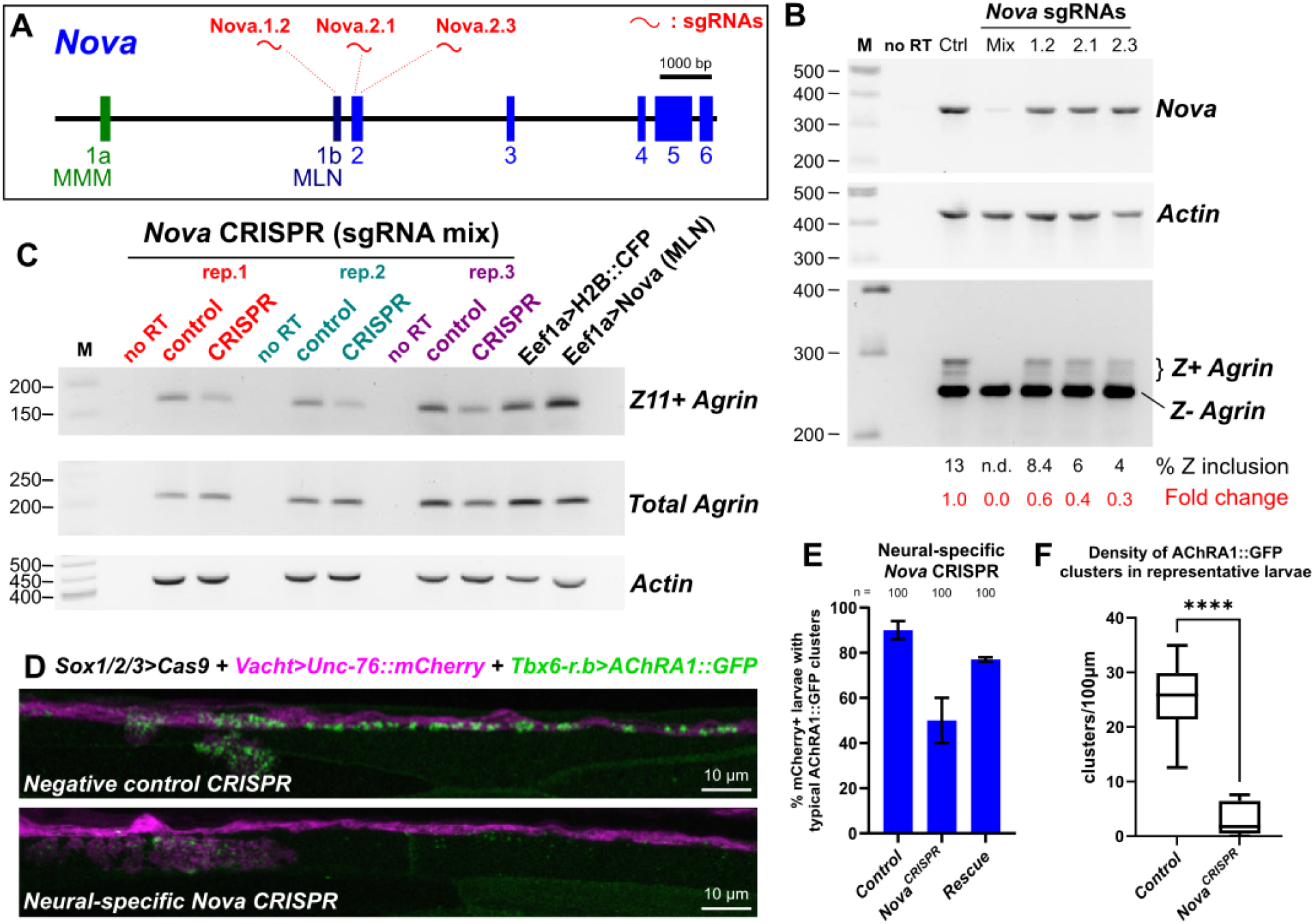
Neural-specific disruption of *Nova* greatly reduced AChR clustering in muscles. A) Diagram of the *Nova* gene in *Ciona robusta,* showing target sites of the three selected *Nova*-targeting sgRNAs. B) RT-PCR assay measuring reduction of *Agrin* Z exon inclusion in larvae upon CRISPR-mediated mutagenesis of *Nova*, compared to a negative CRISPR control sample (“Ctrl”). All three sgRNAs resulted in some reduction of Z exon inclusion on their own, when co-electroporated with neural-specific *Sox1/2/3>Cas9*, but the largest reduction was seen when all three sgRNAs were combined (“Mix”) and co-electroporated with ubiquitously activated *Eef1a>Cas9*. Only the mix substantially reduced *Nova* transcript detection, perhaps due to high frequency of large deletions spanning the target amplicon. M: molecular weight marker. no RT: no reverse transcriptase added. C) All three replicates of ubiquitous *Nova* CRISPR (using *Eef1a>Cas9* and all three sgRNAs combined) show reduction of *Agrin* Z exon band, reproducing the effects seen in panel B. A slight increase in Z exon inclusion is seen upon Nova (MLN isoform) overexpression using the *Eef1a* promoter. D) Neural-specific CRISPR/Cas9-mediated disruption results in decreased AChRA1::GFP clustering in tail muscles, phenocopying neural-specific, CRISPR-mediated disruption of *Agrin* Z exons. Negative control CRISPR performed using *U6>Control* negative control sgRNA plasmid. E) Scoring of larvae represented by the panel above. The loss of AChRA1::GFP clustering was rescued by co-electroporation of a CRISPR-insensitive *Islet -7216/-3950 + bpFOG>Nova(MLN) rescue* plasmid, demonstrating specificity of the CRISPR effect. 50 larvae examined per duplicate and condition. F) Local densities of AChRA1::GFP clusters were quantified in 10 representative larvae in either negative control or *Nova* CRISPR condition (as in panel D) using confocal imaging. **** indicated p<0.0001 following an unpaired T-test using Welch’s correction.

### Expression of *Nova* in neurons is activated by the transcription factor Ebf

Although the role of Nova in regulating neural *Agrin* isoform splicing has been established, almost nothing is known about how *Nova* itself is activated in motor neurons. To help understand the transcriptional regulation of *Ciona Nova,* we isolated a ∼2 kbp sequence immediately 5’ to exon 1b of *Nova,* cloning it upstream of GFP (*Nova[MLN] -2011/+6>GFP,* or simply *Nova>GFP*)(**Figure 7A**). This drove strong GFP expression in MG neurons, in addition to some brain neurons, the otolith, and oral siphon muscle precursors (**Figure 7B**), recapitulating much of the expression observed by ISH (see **Figure 2**).

**Figure 7.**
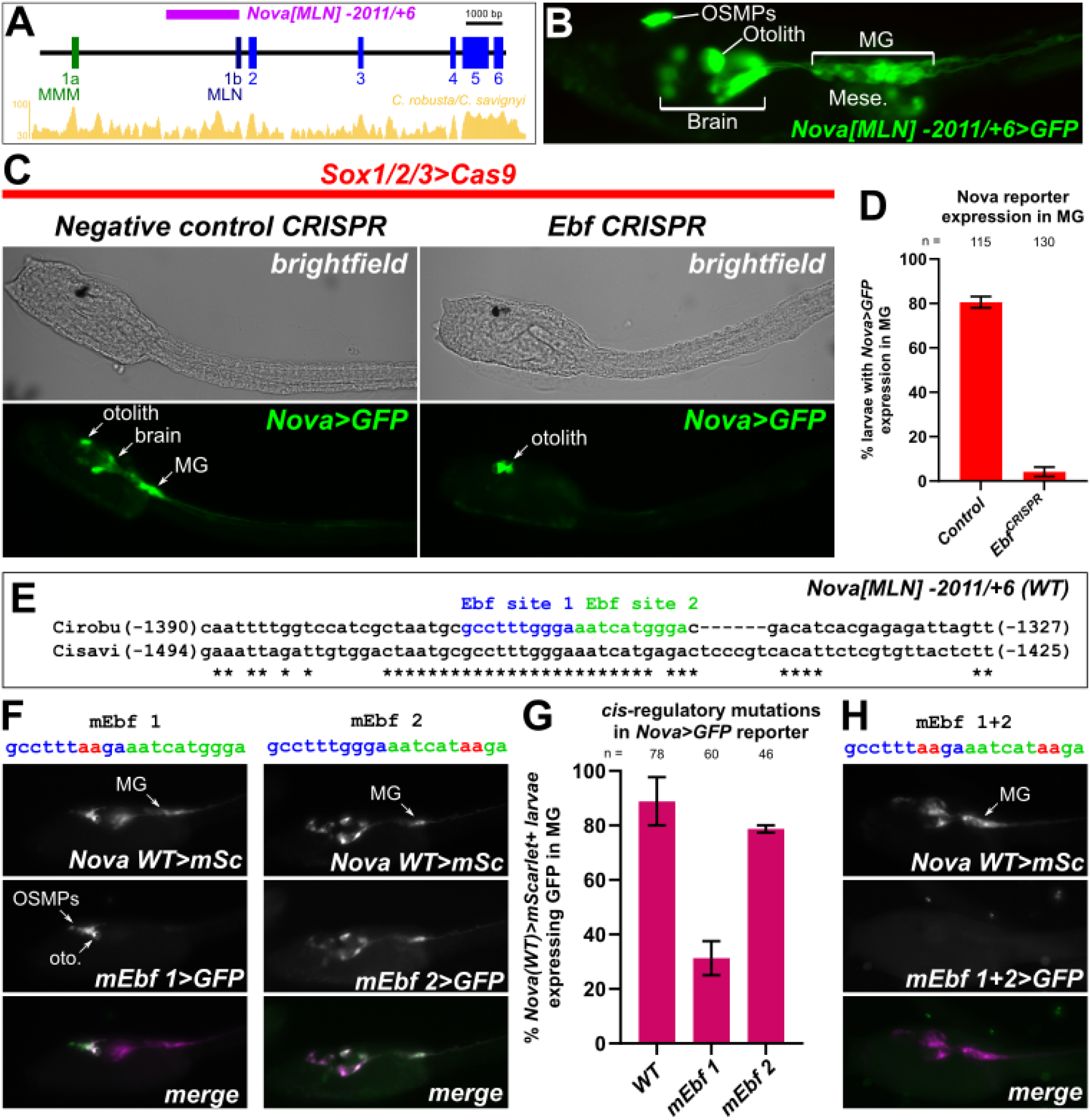
Ebf regulates transcription of *Nova* in *Ciona* motor neurons. A) Diagram of *Nova* locus in *Ciona robusta,* showing position of the cloned *Nova[MLN] - 2011/+6* promoter. Conservation with the *C. savignyi* genome shown as golden peaks below, as visualized in the ANISEED database (Dardaillon et al., 2020). B) *C. robusta* larva electroporated with the *Nova[MLN] -2011/+6>GFP* reporter plasmid, recapitulating expression seen by *in situ* hybridization in the motor ganglion (MG), larval brain/sensory vesicle (including the otolith pigment cell), and oral siphon muscle progenitors (OSMPs). C) Neural-specific CRISPR-mediated disruption of *Ebf* eliminates expression of the *Nova* reporter plasmid in all cells except in the otolith. Negative control performed using *U6>Control* negative control sgRNA instead. D) Scoring of *Nova* reporter expression (as represented in previous panel) in duplicate, with at least 50 larvae examined per duplicate and condition. E) Sequence alignment between *C. robusta* and *C. savignyi* genomic sequences showing conserved, predicted Ebf binding sites ∼1.4 kb upstream of *Nova* exon 1b. F) Effects of *C. robusta Nova* GFP reporter plasmid (green) bearing targeted mutations predicted to disrupt Ebf binding to Ebf sites 1 and 2 (mEbf 1 and mEbf 2, respectively), by co-electroporating the wildtype *Nova* mScarlet (mSc) reporter plasmid (magenta). G) Scoring of larvae represented in previous panel, showing more substantial effect of disrupting predicted Ebf site 1 than Ebf site 2. Electroporations performed and assayed in duplicate, with sample size of at least 15 larvae examined per duplicate per construct. H) Representative image showing complete loss of *Nova* reporter expression upon mutating both predicted Ebf sites.

One of the major sequence-specific transcription factors expressed in the differentiating neurons of the *Ciona* MG is Ebf (Mazet et al., 2005), the sole *Ciona* ortholog of EBF family factors in vertebrates, also known as COE (Collier/Olf/EBF)(Daburon et al., 2008). In *Ciona,* Ebf is expressed in differentiating MG neurons and is required for MN2 specification (Kratsios et al., 2012; Stolfi et al., 2014), while in vertebrates Ebf2 is required specifically for axial MN development (Catela et al., 2019). We therefore sought to investigate the role of Ebf in regulating *Nova* in the *Ciona* MG.

To test if *Ebf* is required for *Nova* expression in differentiating MG neurons in *Ciona*, we targeted it for neural tissue-specific CRISPR/Cas9-mediated disruption using a previously published, highly efficient sgRNA (Gandhi et al., 2017). We assayed loss of *Nova>GFP* reporter gene expression in the MG neurons of *Ebf* CRISPR larvae compared to control larvae. Neural-specific disruption of *Ebf* (using again *Sox1/2/3>Cas9*) resulted in significant, near total loss of *Nova>GFP* in MG neurons (**Figure 7C,D**). Taken together, these data suggest that Ebf is necessary for neuron-specific expression of *Nova* in *Ciona* embryos. Curiously, *Nova>GFP* was not lost from the otolith, which does not express *Ebf*. This suggested the possibility of Ebf- independent regulation of *Nova* in this cell.

To get a better understanding of whether *Nova* is a direct or indirect transcriptional target of Ebf, we searched the *Nova* 5’ *cis*-regulatory sequence for potential Ebf binding sites. The predictive algorithm JASPAR only found two such sites, in tandem ∼1300 bp upstream of the transcription start site and almost perfectly conserved in the related species *C. savignyi* (**Figure 7E**). When we mutated the 1^st^ Ebf site (mEbf 1), reporter expression was greatly reduced in MG and brain neurons, but not in the oral siphon muscle precursors or otolith (**Figure 7F,G**). In contrast, when we mutated the 2^nd^ Ebf site (mEbf 2), reporter expression was not significantly reduced (**Figure 7F,G**).

However, when both sites were mutated in the same construct, we lost all expression in Ebf+ cells (**Figure 7H**), suggesting that the Ebf 2 site might serve as a “backup” site for the primary site, Ebf 1. Taken together, we conclude that Ebf is a key activator of *Nova* transcription during motor neuron differentiation in *Ciona*.

## Discussion

Here we have shown that a conserved alternative splicing-based switch for acetylcholine receptor clustering at neuromuscular synapses is shared by vertebrates and their close relatives the tunicates (**Figure 8A**). In this evolutionarily conserved pathway, the RNA-binding protein Nova is expressed in motor neurons and promotes the inclusion of *Agrin* “Z” microexons, which encode a short peptide motif that mediates the activation of Lrp4 and downstream clustering of AChRs post-synaptically in muscle cells. To our knowledge, this is the first report of a Nova-Agrin-Lrp4 pathway for AChR clustering outside the vertebrates, including non-vertebrate chordates such as amphioxus (Irimia et al., 2011), pushing the evolutionary emergence of this mechanism at least as far back as the last common ancestor of tunicates and vertebrates.

**Figure 8.**
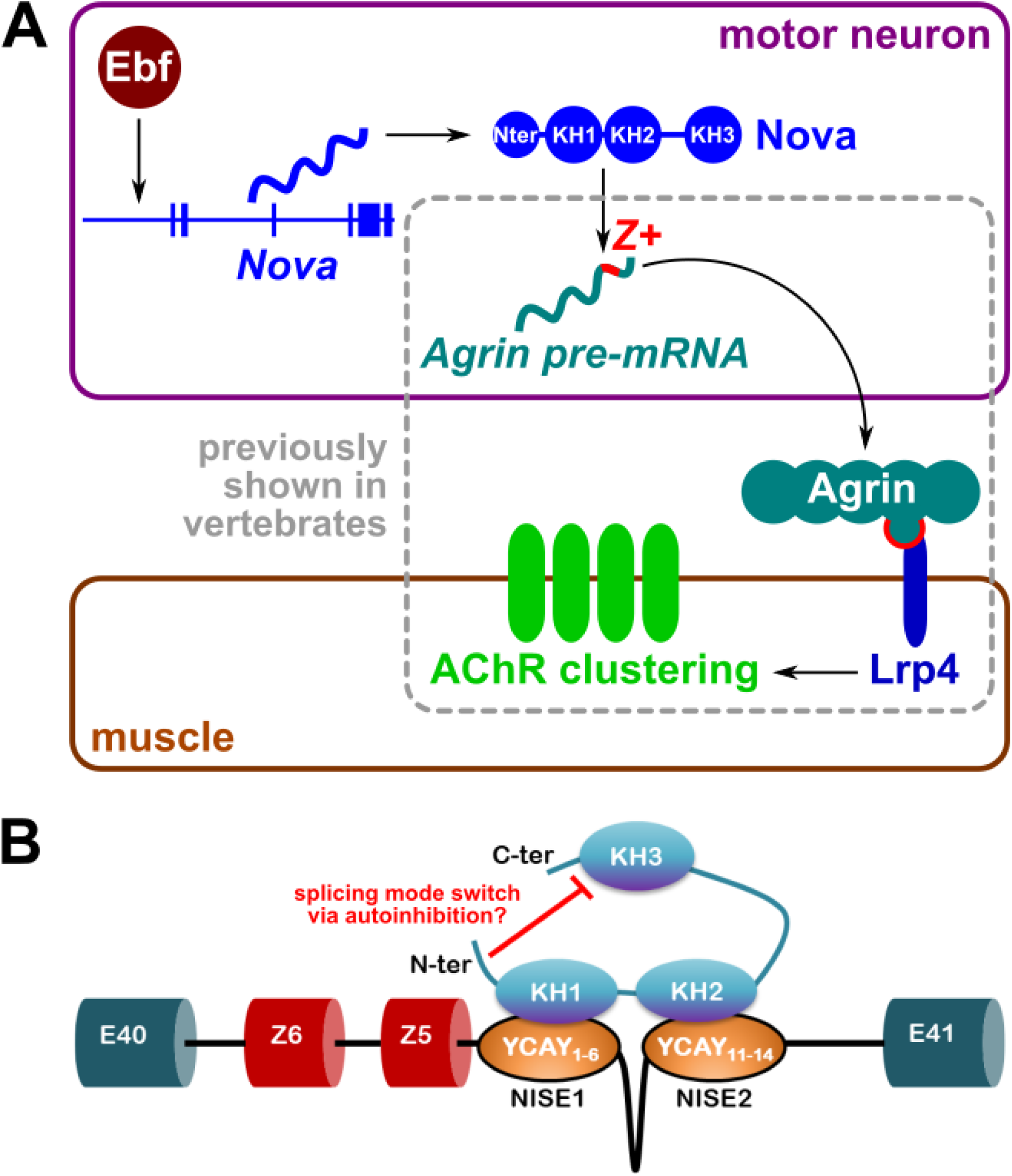
Summary model diagram of a conserved *Nova-Agrin-Lrp4* pathway for AChR clustering in *Ciona.* A) Summary of conserved pathway, and previously unidentified Ebf-Nova regulatory connection identified in *Ciona.* B) Model of proposed mechanism for *Ciona* Nova- dependent alternative *Agrin* splicing via binding of KH1 and KH2 domains to two YCAY- rich NISEs identified in the intron 3’ to the Z exons. Inhibitory effect of KH3 domain is relieved by the N-terminus. See text for details.

Although we have shown that the basic regulation of *Agrin* Z exon inclusion by Nova is deeply conserved, it is not yet known which KH domains of Nova1/2 mediate Z exon inclusion in vertebrates. It is also not known which *cis-*acting sequences might mediate Nova1/2 binding to vertebrate *Agrin* mRNAs. Although the general pathway is conserved, the exact mechanisms of *Agrin* splicing by Nova might be divergent, as mouse Nova1/2 proteins were unable to promote Z exon inclusion in a *Ciona Agrin* minigene assay. Using the same assay, we revealed a potential autoinhibitory mechanism involving the N-terminus and the KH3 domain of *Ciona* Nova, which could either be conserved in vertebrates, or might explain their apparently divergent mechanisms of Nova splicing activity. Given that *Ciona* Nova needs both KH1 and KH2 domains to promote Z exon inclusion, and this depends on two separate NISEs in *Ciona Agrin*, we propose a model where KH1 binds to the first NISE and KH2 to the second NISE, or vice-versa. Meanwhile, KH3 may be available for interactions with pre-mRNAs encoded by other genes, unless inhibited by the N-terminal domain (**Figure 8B**). Further work in both *Ciona* and vertebrates will be required to investigate in greater detail the evolution of Nova structure-function.

Finally, we have also shown that the transcription factor Ebf, an important terminal selector of motor neuron fate (Kratsios et al., 2012), is required to activate transcription of *Ciona* Nova. Ebf orthologs are also expressed in vertebrate motor neurons (Catela et al., 2019), suggesting the possibility that the regulatory connection between Ebf and *Nova* might also be conserved. This would in turn connect a largely RNA- and protein-based pathway for AChR clustering (Nova-Agrin-Lrp4) back to a transcriptional gene regulatory network downstream of motor neuron specification.

## Materials and methods

### *Ciona* handling and electroporation

Adult *Ciona robusta* (*intestinalis Type A*) specimens were collected and shipped by M-REP (San Diego, California) and kept in artificial sea water tanks until use. Gametes were isolated and dechorionated as previously described (Christiaen et al., 2009c).

Electroporations were performed as previously described (Christiaen et al., 2009b), using plasmid DNA mixes defined in the **Supplemental Sequences File.**

AChRA1::GFP plasmid (Nishino et al., 2011) was kindly provided as a gift by Dr. Atsuo Nishino. For direct visualization of GFP/mCherry fluorescence, embryos and larvae were fixed in MEM-Formaldehyde buffer as previously described (Johnson et al., 2024). Fluorescence whole-mount mRNA *in situ* hybridizations were performed as previously described (Ikuta and Saiga, 2007). Embryos and larvae were imaged using upright or inverted epifluorescence or scanning-point confocal microscopes.

### CRISPR/Cas9 methods in *Ciona*

Internet-based prediction algorithm CRISPOR (Concordet and Haeussler, 2018)(http://crispor.tefor.net/) was used to identify candidate sgRNAs for CRISPR/Cas9. Expression plasmids for sgRNAs were constructed by ligating annealed oligonucleotides (Stolfi et al., 2014), Gibson assembly of PCR products (Gandhi et al., 2018), or synthesized and custom-cloned *de novo* (Twist Bioscience, California).

Validation of sgRNA efficacies was performed using either Sanger sequencing of amplicons following the “peakshift” method (Gandhi et al., 2018), or by Illumina-based next-generation amplicon sequencing (Johnson et al., 2023). All promoter, sgRNA, and primer sequences listed in the **Supplemental Sequences File.**

### Minigene assay

The day before the transfection, 0.6 x 10^6^ HEK293T cells were seeded per well in a 6-well plate (USA Scientific) in DMEM culture medium. On the day of transfection, a total of 2.5 µg DNA of minigene, cDNA construct, and empty vector was used to transfect each of 6 well plate(s) and 7.5 µL of linear polyethylenimine (PEI; Polysciences), MW 25,000 (1mg/mL) was used in a ratio of 1:3 (DNA : PEI). 0.5 µg (= 1x) of minigene DNA was used in each well to test splicing with different amount of splicing factor (0 µg = 0x, 0.5 µg = 1x, and 2.0 µg = 4x). Empty vector was used to bring the total amount of DNA to 2.5 µg (2.0 µg = 4x, 1.5 µg = 3x, and 0 µg = 0x) per well.

The total volume of the DNA mixture was 200 µL (Table 6 and 7). First, the exact amount of DNA in µL was pipetted in a 1.5 µL Eppendorf tube (Eppendorf) and Opti-MEM media (Thermo Scientific) was used to bring the volume to 192.5 µL. Then the mixture was vortexed thoroughly. Finally, 7.5 µL of PEI was added, vortexed, and centrifuged briefly. The mixture was then incubated for 15 minutes at room temperature.

In the meantime, medium in the cells was aspirated and 2 mL of fresh DMEM medium was added. After a 15 minutes incubation, 200 µL of reaction mixture was added to the cell and the plate was cross-shaken gently. The plate was then incubated for 48 hours at 37°C. All cloning primers listed in the **Supplemental Sequences File.**

### RT-PCR and qPCR

RNA from homogenized HEK293T cells was extracted 48 hours after transfection using RiboZol RNA Extraction Reagent (AMRESCO) or IBI Isolate (IBI Scientific) according to the manufacturer’s instructions. A total of 5 µg of RNA per sample was digested in a 50 µL reaction containing 1.5 µL of TurboDNase (Thermo Scientific), 5 µL of 10X Buffer, and double-distilled water (ddH2O) to 50 µL. After 30 minutes of incubation at 37°C another 1.5 µL of TurboDNase was added to the mixture and incubated for another 30 minutes. After a total of one-hour incubation 10 µL of TurboDNAse Inactivation Reagent was added and samples were kept at room temperature for 5 minutes, and the tubes were flicked every 2 minutes to resuspend the inactivation reagent. Then the tubes were centrifuged at 10,000 rpm for 90 seconds to collect supernatant for processing.

From total RNA we synthesized cDNA using RevertAid First Strand cDNA Synthesis Kit (Thermo Scientific). A mix of 250 ng RNA and 1 μL of oligo (dT)18 at 500 ng/μL was prepared in a total volume of 12 μL (diluted in sterile ddH2O). The mix was incubated for 5 minutes at 65°C in a PCR machine. After this, an RT reaction mix was prepared combining the mixture above with the following ingredients in a total volume of 20 μL: 5X RT Buffer (4 μL); RiboLock RNase Inhibitor 20 U/μl (0.5 μL); 10 mM dNTPs (2 μL); RevertAid RT 200 U/μl (0.5 μl). This mixture was incubated for 1 hour at 42°C followed by 5 minutes at 72°C in a PCR machine. After the incubation, 5 μL of water were added to each tube bringing the volume to a total of 25 μL. Each RT reaction mix had a concentration of 10 ng of starting RNA/μL. 5 μL from each RT reaction, equivalent to 50 ng of starting RNA, were used as template for each RT-PCR.

A mixture of 10X PCR Buffer (5 μL); dNTPs 10 mM (1 μL); forward and reverse primers each 10 μM (1 μL); 5 U/μL HotStarTaq Plus DNA polymerase (Qiagen) (0.4 μL) or 5 U/μL Dream Taq Hot Start DNA polymerase (Thermo Scientific) (0.4 μL) and RT reaction (5 μL) in a total volume of 50 μL (diluted in sterile ddH2O) was prepared in a PCR tube. The PCR reaction was performed with initial denaturation for 5 minutes at 95°C; variable number of cycles of: denaturation for 30 seconds at 94°C, annealing for 30 seconds at 60°C and elongation for 30 seconds at 72°C. This was followed by a final extension of 7 minutes at 72°C and a hold at 12°C. All primer sequences can be found in the **Supplemental Sequences** file.

## Supporting information

Supplemental Sequences File

## Acknowledgments

We are grateful to Atsuo Nishino for sharing the AChRA1::GFP construct. We thank members of our labs for critical feedback and suggestions. We thank Susanne Gibboney for technical assistance. This work was supported by grants R01HD104825 from NIH/NICHD and 1940743 from NSF/IOS to AS, grants R01GM96032, R01HL108643, and R01HD096770 from NIH to LC, and grant R15GM119099-01 from NIH/NIGMS to MR.

**The authors declare no conflicts of interest**

---

**Figure S1.**
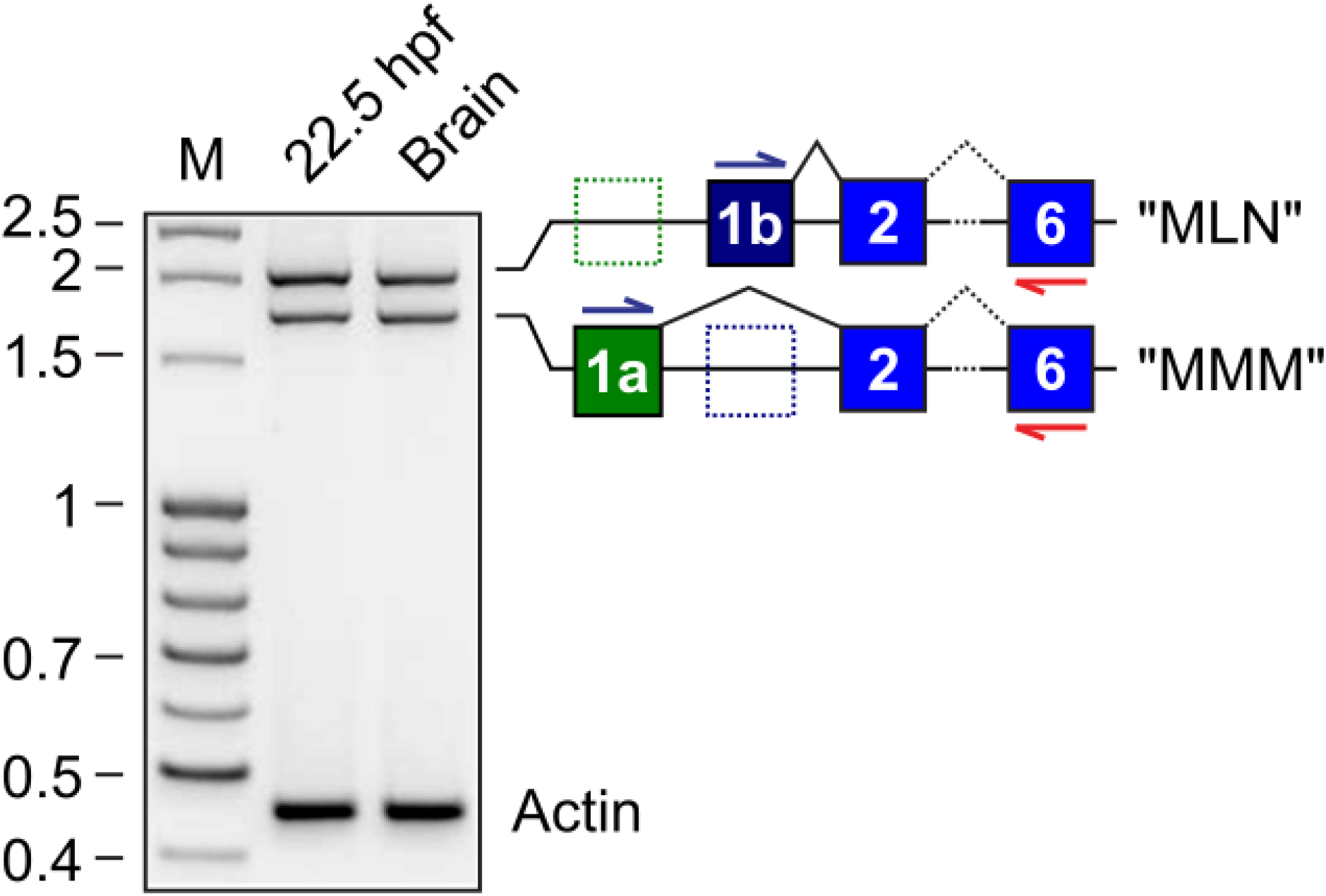
RT-PCR detection of different *Nova* alternative splice forms in larvae (22.5 hours post-fertilization) and adult brain. M: DNA molecular weight marker.

**Figure S2.**
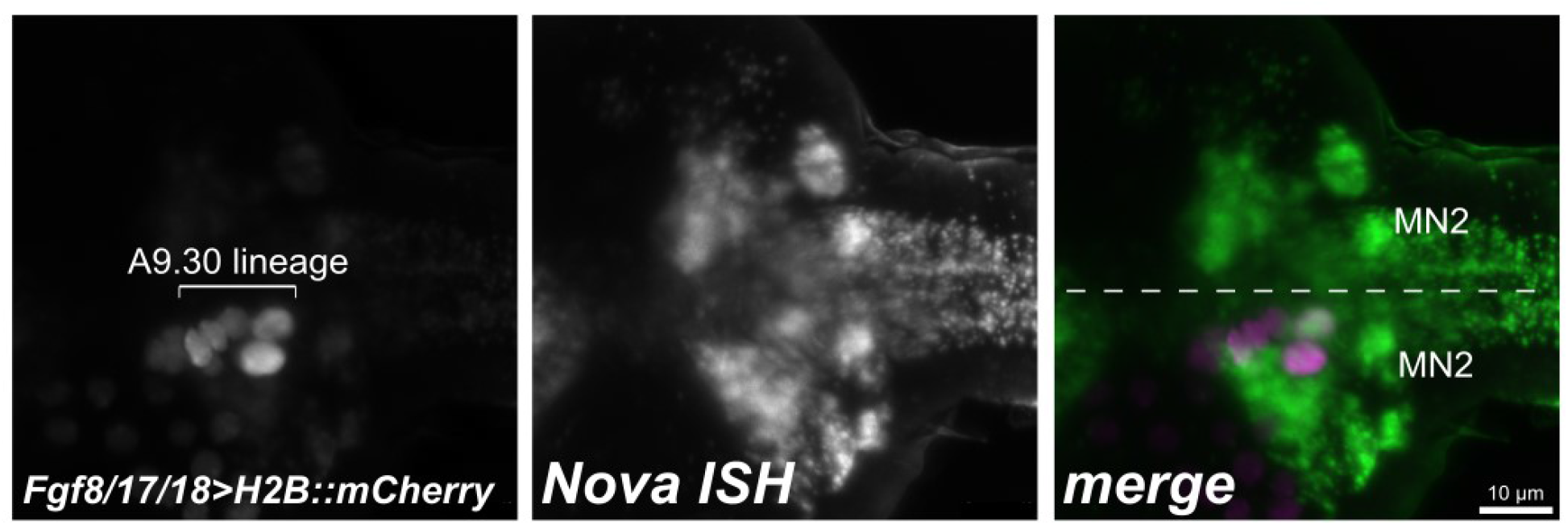
*Nova* mRNA *in situ* hybridization coupled to immunostaining-based detection of Fgf8/17/18>H2B::mCherry expressin, revealing identity of Nova+ MN2 cells adjacent to the *Fgf8/17/18* reporter-expressing cells.

**Figure S3.**
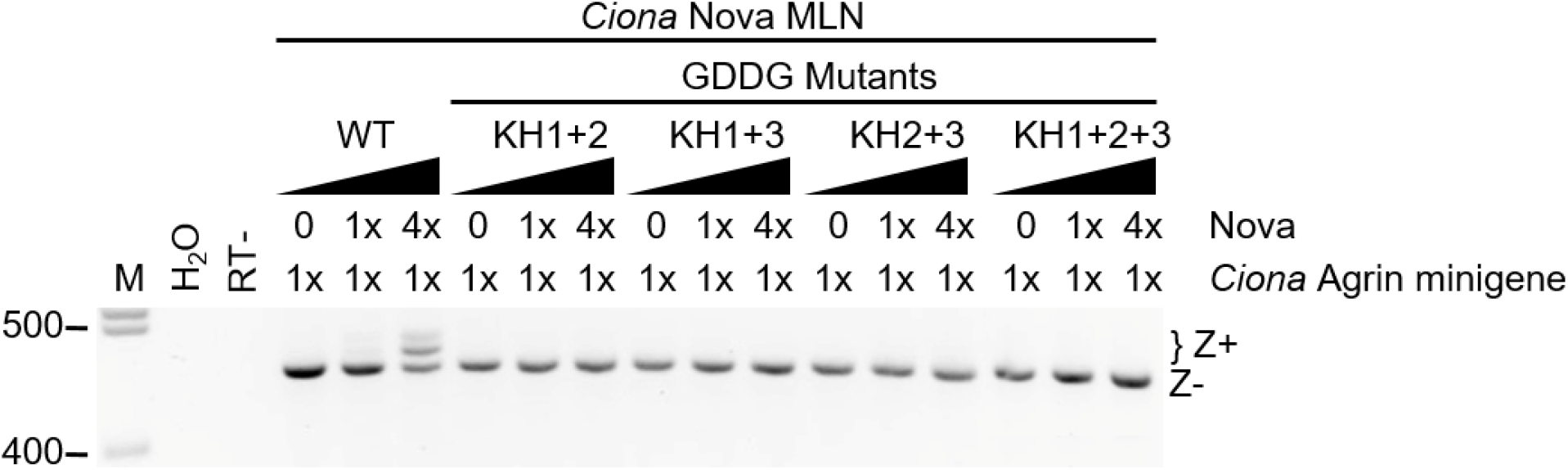
*Ciona Agrin* minigene assay showing effect of combining different CDDG mutant KH domains. M: DNA molecular weight marker. H2O: using water instead of cDNA template for PCR. RT-: no reverse transcriptase added.

**Figure S4.**
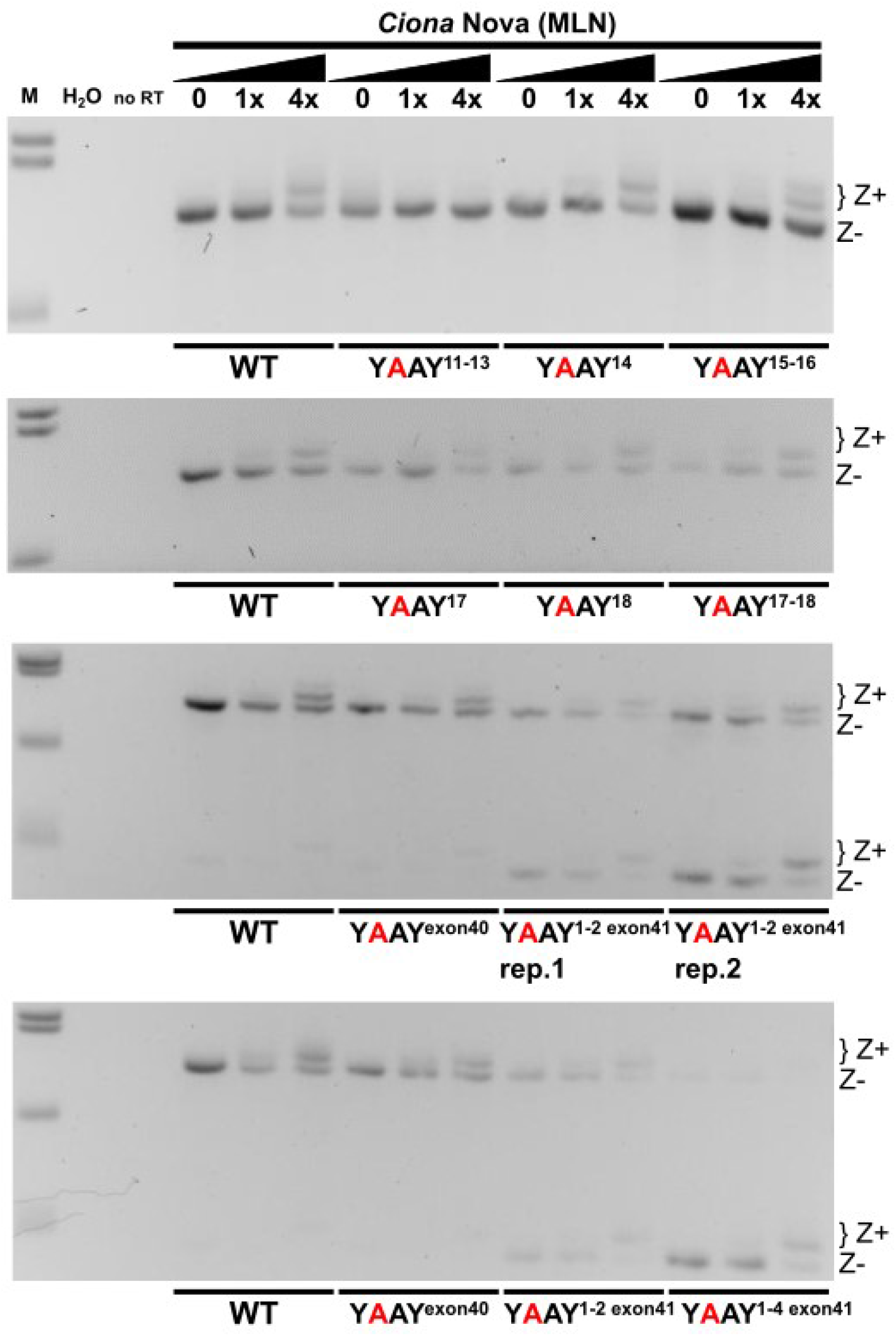
Additional candidate YCAY site mutagenesis experiments. Experiment performed, assayed, and presented as in main figure 4. Smaller products seen with exonic YCAY mutations likely represent aberrantly-running products. M: DNA molecular weight marker. H2O: using water instead of cDNA template for PCR. no RT: no reverse transcriptase added.

**Figure S5.**
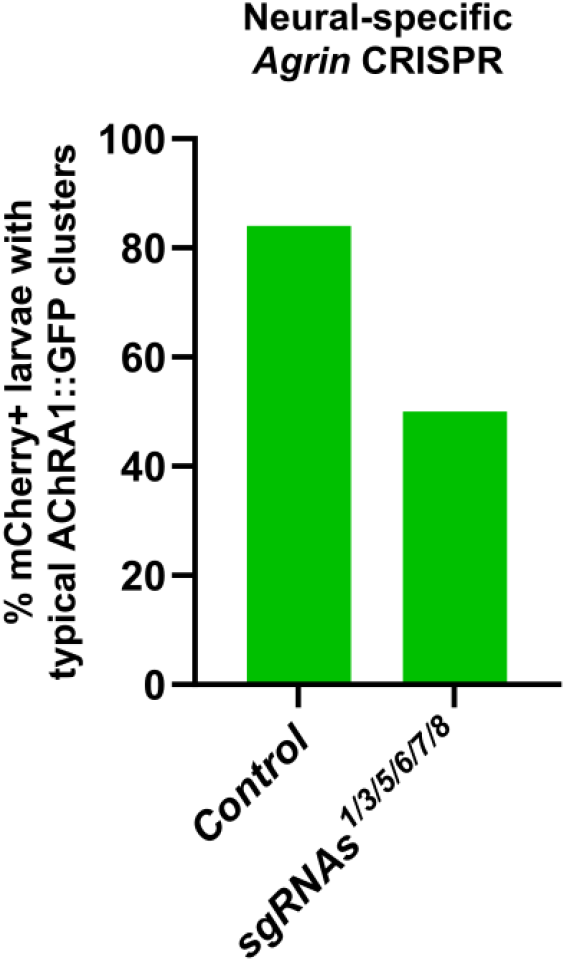
Scoring of AChRA1::GFP clustering upon initial neural-specific CRISPR-based disruption of *Agrin* using 6 different sgRNA expression cassettes in the form of PCR products.

**Figure S6.**
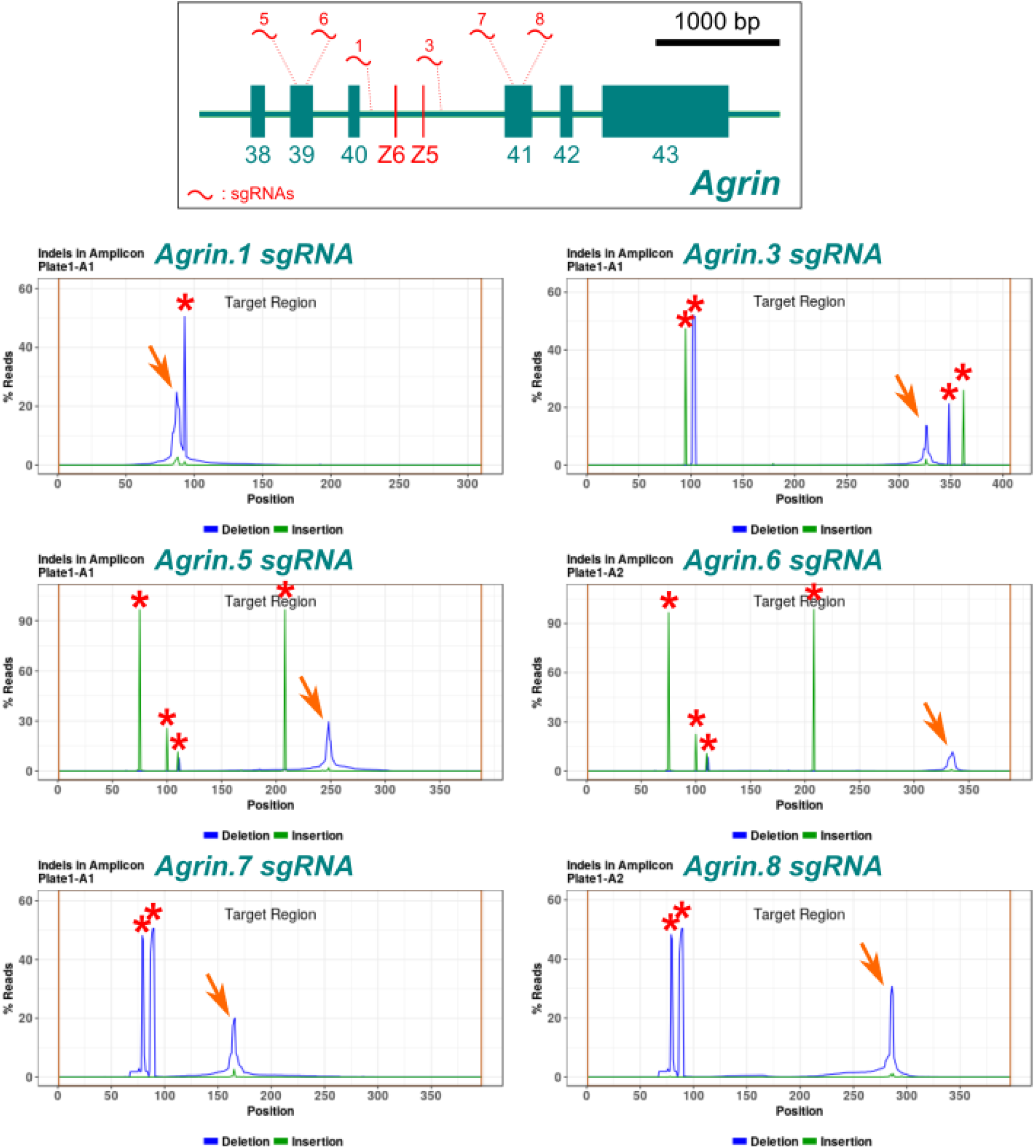
Illumina sequencing-based validation of selected *Agrin* sgRNAs. Red arrows indicate indel plots showing estimated sgRNA efficacy. Red asterisks indicate naturally-occurring indels. *Agrin* sgRNAs 2 and 4 were not validated due to their inclusion in the seemingly least effective sgRNA combination, combo #2 (see Figure 5).

**Figure S7.**
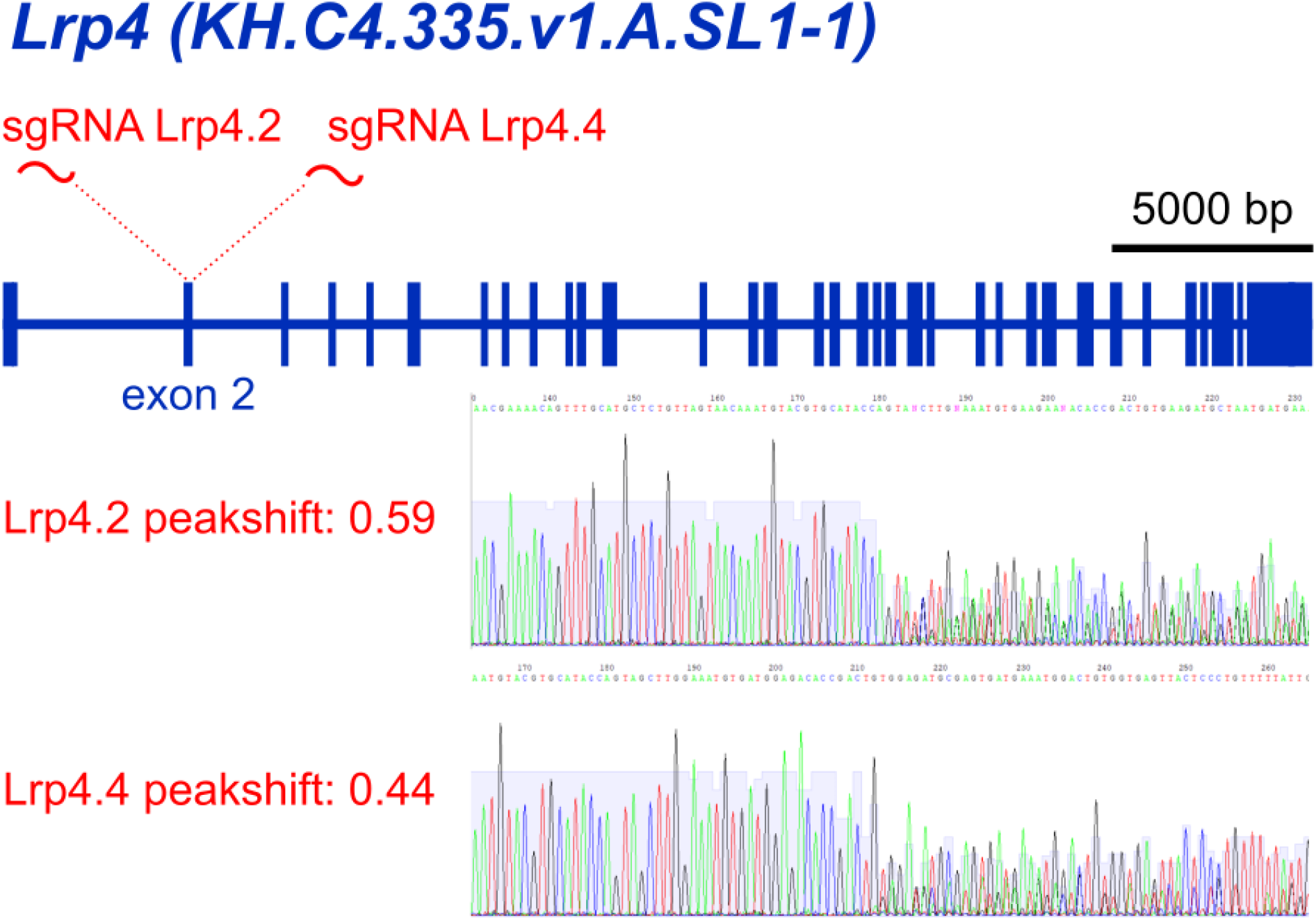
“Peakshift” (Sanger sequencing-based) validation of *Lrp4*-targeting sgRNAs.

**Figure S8.**
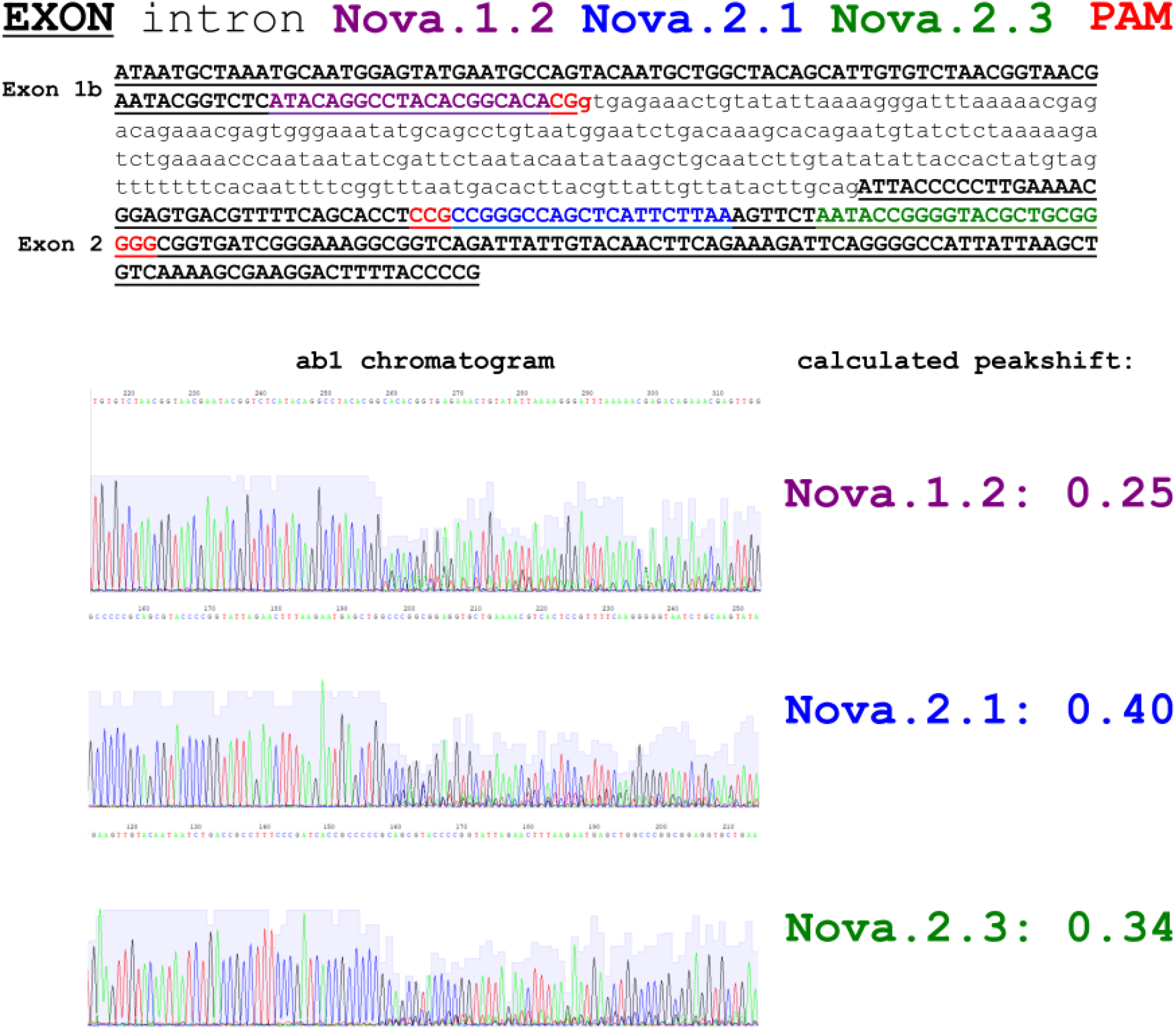
“Peakshift” validation of *Nova*-targeting sgRNAs.

## Notes

### Competing Interest Statement

The authors have declared no competing interest.

