## Supplemental Sequences File for "A conserved RNA switch for acetylcholine receptor clustering at neuromuscular junctions in chordates"

**Supplemental Sequences Files**

**Published *in situ* probe templates:**

*Islet -* Cirobu.g00011396 (Satou et al. 2002)

**New *in situ* probe templates:**

*Nova (C. robusta)* *in situ* probe template

ATGGAGTATGAATGCCAGTACAATGCTGGCTACAGCATTGTGTCTAACGGTAACGAATACGGTCTCATACAGGCCTACACGGCACACGATTACCCCCTTGAAAACGGAGTGACGTTTTCAGCACCTCCGCCGGGCCAGCTCATTCTTAAAGTTCTAATACCGGGGTACGCTGCGGGGGCGGTGATCGGGAAAGGCGGTCAGATTATTGTACAACTTCAGAAAGATTCAGGGGCCATTATTAAGCTGTCAAAAGCGAAGGACTTTTACCCCGGAACCCAAGACCGAGTCGTTTTGATCCAAGGAACCGCCGAAGGCTTGATGAAGGTGCAAAATACCATTATAGAGAAGGTGTACGAGTTCCCTGTGCCCAAAGATTTAGCTGCGATCATCGGAGACCGACCGAAACAGGTGAAAATCATCGTACCCAACACAACTGCGGGACTGGTAATAGGAAAGGCCGGCGCAACGATAAAGACCATTATGGAAGAGAGTGGATCGAAGGTTCAACTCTCGCAAAAGCCAGACGGGGTAAACGTCCAAGAACGAGTCATCACAATCAAAGGAGAGAAGCACCAACTCATGACAGCATCTAATATTATTATTGATAAAATTAAAGACGACCCTCAAAGCGCCAGTTGCCCTCACATAAGTTACTCTGGCATCGCTGGCCCGATCGCTAACGCGAATCCCACCGGATCGCCCTACGCTGCTGGCTCGGCTGCATTAGTTGACGCTTCGCACCCATCCGTGGCCGCTATGTTGGGACATTATGTTATCCCAGGCCAACAGGTGCTGCAGACAGCAATGCCACTCTCCCATCACCCGCACCAGTCCGCGTTGTCCAGCGGCTCAGTGACACCGGCGCCTGAACTGACGACCATAAACCACGCCATGACAACGTTAGCGAACTATGGCTACACCTTAGGAGGCGTAAACTATGGTACCTTGGGTGTAATGCCTAGTGTACATCCAAGTGTACACCCTGGCATCGCTACCTCGGTCGGGATGATCTCTGCAGGCTCCCTAGCAGGAAGTCCAATCCCTTCAGCTACCCCCTTGCTCTCTGCCACTGCTCTACCGACGGAATCCAGTATTCCGACGGCTGTTCCCACTGCCCAAGCCATTTCAATGCAGAGCAATTACCTTGCAAACTTGGCTAATGCTGGTTACCTGACTACCGGTCACCCACAGTTGCTTGGAGCGACGTCAGGCCTCGGCGGTCTCACCACAGTGTCCCAGCACCCGCCACCAGCGGCGACACCAACGAGTTTTTCCGTCGCTTCTACCCCTTCTACCCCTGGTCTGCCGGTTTCATTTAGCCCCCATTCAACCGTGAGTATCCTAAGCATCGAAAAGTCAAGCGACGGACAAAAAGAAACAATTGAACTGGCAATTCCCGAAAACCTCATCGGAGCAGTCCTCGGAAAAGCGGGAAGGACACTGGTTGAGTATCAGGATGTATCAGGGGCGAAAATTCAAATTTCTAAAAAGGGTGATTACGTCGCCGGGACCAGGAACAGGAGGGTTACGATTACGGGGAAGCCCCCATGCCCACAGACTGCGCAGTTTCTTATTACGCAACGTGTCGCCTCTGCGCAGAACGCAAGGGCACAGCAGGCTAAGTTACTGTAGGTCAGGAACCACGCCAGTCTTCCACAAATGTGCTGCCGCATATTGTTTTACCTTAGTAGTTTGTACTTTTAGTAAGGTTGAATTTTTCGCACTGAGGAGCGCTAAGTACGTTTCTTAAAAAGTCTCTTCGCCAAATCAAGTCTTCCGGGCCATTCGCCAAAATCGCATCTAAATTCGATCATCTTAAACAGCGTACACGACTTTGCTTTTTTCGGCGTTCACGAAACGCGCTCATACATTATTCAAGTCAATTACGAAATCTCTGCGAGTTTGTGACGCACTACTGTGCACTACCATCGATCCCGCTTTTCCGTAT

*Agrin (C. robusta)* *in situ* probe template

CGTTGGTTTCAGTGCGTTTAGGATTGAGTTTCGCTTCTTAAAGAGAAAAATACAGCGCAAAAAAAGATAAAACAATACAGTGACGGGGTAAAAACTAAGAAGCGAAAATTTGTTCGGTCCCCAGGAAAATAAAAAATAAATAGAGCATGCTACGGTATGGTACAACATGTAGAATTATCTAAAAAGTTCAAATAAGAATATGGAATGAAAGGAGACCGAGAGAAATCGACAGTACGTGCGGTATACACCCTCGCAAAAAATCAAAACTTTTTTGTCGTTTTTAAAATCGAAAATTATAAATGTAAAGGCTTCGTAACCTTAATAAAGTGATTGTTCGAAACGATGCAAAACACGAAAGTGTGAAAATTGAAATAAAATGCTAAAGGTGAAAATATGGTAAATGTATTGAAATCTGATACCCATGCTTAGACAACAGTAAAAACTAAGCGCCTAAATTTGGTGGCAATATTGTAAAGAAATTTTGCGCCAGGTATTTAACCAGTTCAAATGAAAATGTATTCACTTGTTACTATATTAGAAAATTTAAAAGACATTTTGTTTTTTTTCATGGTGATGCAAAATTGAAAAATTACCTAAAATGATTTCCGTTTCGAATACGTTAGTGGGTGTGGATAATAAAGTGTGAAAGAAATTGTTTTGATGGTTGTGCTGTGCTCGAACCTACGGCGCTGTCAGCTTTAAAATCTGTGGCTTGCGTGTATCCATCATCGACCATTACACGAACGCACAACGGGAGAATTAAGCGCGTCAGCCCGTAAATCAATTGATTCGTCGTGGATCTTTGCTGAGGCCACGCAGCCGACGAACGGGACAAAGAAAGATTTGTAAAGACCAATACCTCGCGGTCGGTAGTCAACGCCACCTAACCACAACATTCCATCAGAGTTCAGGTGATTAGTTAACCCTGGTGAGGTCGCTGTCTTTATTAACCCGTTGTTTACTTGCAAACTTCCTATGTTCATTTTCCTGTTAACTTTTACAGTTGTCCATTCTCCGTTGTTGATTTGCTGGTCGGATATTACGTGCGCTGGACCAGAACCAAGGTCAAAACGAAGATGAAGACGACCGTCATGGATTGCGAGCGCAATATAATCGACCCCCTCCCTCGCTTTGCCCACCATGAGCAGTAGACCGTGACGAGCAGTAGTGCGGAACACGATTTCATAATTGTTGTGTGTTCGTGCACGAGAATTGTCACTGAGTTGCTCTCGAAAATCGTTTGGCATTGCTTTCACTGCATTCCTATACATTATCTTAGTTGTTCCATCCAAATAAATGGCAGTCGCTTGTTCATCTTGTAGCAAGTCAGTAGAATGTTCTTGCTCACAGTTATCTCCCGTGTAATAAGGTAAGCACACGCACATATATTCTGCTCCACGTGGATGGCATACACCACCATTATCGCATGGGTTGCGATAACATGTATGGGCGTTGAATGCAATTACATTGAAGTAAGAGACACTACCAGCTGGGGAAATGGGCAGGTTCACACCGTTCACTTGGAACTTCTGTAATGCACCACTCAGTCCAGTTGTCACTCCAGCTTCTGGATTAAATTTGACACCATCAGGGAACCCACCAACATACATAGGTTGCTTAAGGTCTAAGAAGGAGTGTTGGCTCGGTGATGTCCCATACACTGGATCAAAATTATCCAATGAAAGGTCTCCAGTTCTCATGGCACGAGATAACACCACAATATGCCATTCATTCAAACTTACAGGGTTGGCACTCCTGATATTTGCTGCCCCTTGTCCAAGGTTGTATTTAAACTCAAGGAATCCATTCTTTAGATTGAGGGAAACAAAATCTCCTTTGCCAGATTTTTTCTGCCCGTTGTAGAAAATTAAACCGTCAGGTTGATTTGAATAAAATAAGATTTCTATTGACATGATTGACCGAACATCCTTTCCTAAAGATGGAAGTTCCAAATATGAATCGCCGGCAAATGCAG

**Validated sgRNAs used in this study (N19):**

Nova1.2: TACAGGCCTACACGGCACA (Vitrinel et al. 2023)

Nova2.1: TAAGAATGAGCTGGCCCGG (Vitrinel et al. 2023)

Nova2.3: ATACCGGGGTACGCTGCGG (Vitrinel et al. 2023)

Agrin.1: AGGTGGCTTGTACTAACAG

Agrin.3: GTCCATAGGAATGTGCGCA

Agrin.5: TAAGAGACACTACCAGCTG

Agrin.6: GTGGTGTATGCCATCCACG

Agrin.7: CACGGTCTACTGCTCATGG

Agrin.8: CCGACCAGCAAATCAACAA

Lrp4.2: TACGTGCATACCAGTAGCT

Lrp4.4: TCACTCGCATCTCCACAGT

Ebf.C: AGACTGTGCCAAGACACCC (Gandhi et al. 2017)

**Negative control sgRNAs:**

Control: CTTTGCTACGATCTACATT (Stolfi et al. 2014)

DenhT2: CGAAATTGCTCGACGCGCT (alternate negative control sgRNA)

**Other sgRNAs designed but not used or validated:**

Nova1.1: GCTACAGCATTGTGTCTAACGG (no discernible peakshift)

Nova2.2: CTAATACCGGGGTACGCTGCGG (no discernible peakshift)

Nova2.4: CAACTTCAGAAAGATTCAGGGG (no discernible peakshift)

Nova3.1: TGATCCAAGGAACCGCCGAAGG (no discernible peakshift)

Nova3.2: ACCTTCATCAAGCCTTCGGCGG (no discernible peakshift)

Lrp4.1: GCCAAGCAGAAAATAATCCAGG (no discernible peakshift)

Lrp4.3: CAGTAGCTTGGAAATGTGATGG (peakshift 0.43)

Lrp4.5: TTGTCGCATTGAAACGAAGAGG (peakshift 0.04)

Lrp4.6: TGGCAGAATGGAAATGCGATGG (peakshift 0.15)

Agrin.2: GTGGCTTGTACTAACAGTGGGG (untested)

Agrin.4: TCCATAGGAATGTGCGCACGGG (untested)

**Peakshift PCR primers used in this study:**

Nova Exons 1b+2 Forward: TCTATGAACTGAGGTCGCAAG

Nova Exons 1b+2 Reverse: CACCTACCTAAATGGACCAAC

Lrp4 Exon 2 Forward: GTGATGTTGTCGTTTCTCATGC

Lrp4 Exon 2 Reverse: TGTTCGCACTTATTTGTAAACCG

**NGS PCR primers used in this study:**

Agrin.1+2.NGS.F: TCTAGCACAAAAGCATATCAC

Agrin.1+2.NGS.R: TCTGCTTCACAAAATTTCAGT

Agrin.3+4.NGS.F: TTCGAGAGTAAGTGGTTATGC

Agrin.3+4.NGS.R: TTTCATACAGCACTATTGACA

Agrin.5+6.NGS.F: CAACTGGACTGAGTGGTGCAT

Agrin.5+6.NGS.R: TAAGGTCTCACCTTGCTCACA

Agrin.7+8.NGS.F: GAAGCACACACACCCTAAATC

Agrin.7+8.NGS.R: TTATAGCAGGGTTTAAGCCGA

**RT-PCR primers used:**

Z11 Agrin isoform:

Ciona Agrin Z6+Z5 Forward 1: TTCGAGAGCAACTAAGTGACAA

Ciona Agrin exon41 Reverse 1: ACGAAGATGAAGACGACCGT

All Agrin isoforms:

Ciona Agrin exon 40 Forward 2: CAAGATGAACAAGCGACTGCCATT

Ciona Agrin exon 41 Reverse 2: GATATTACGTGCGCTGGACCAGAA

Minigene assay primers:

pCI Agrin RT F: GTGTCCACTCCCAGTTCAATTACAG

pCI Agrin RT R: TGTCTGCTCGAAGCATTAACCC

**Other published plasmids:**

*Fgf8/17/18>H2B::mCherry* (Stolfi and Levine 2011)

*Islet -7216/-3950 + bpFOG>Unc-76::mCherry* (Stolfi et al. 2010)

*Eef1a>Cas9* (Stolfi et al. 2014)

*Sox1/2/3>Cas9* (Stolfi et al. 2014)

*VAChT -4315/+15>Unc-76::mCherry* (Popsuj et al. 2021)

**New construct based on new combinations of published sequences:**

*Tbx6-related.b>AChRA1::GFP*

Tbx6-related.b driver (Christiaen et al. 2009)

AChRA1::GFP (Nishino et al. 2011)

ggcgcgcccaacggagtacgcgtgtcaagtttaatggcgtaattaccgaacaactgttgataagtaatgaggaccccgctgcggtaacctttcgaaattgctggttgcaagcggtgttggtgcgataataaggaaaagcggtaagcgttattttttgatccaccacaaagaccaaacctatattgatacactaaactaaatcaaattcgaacctatccattaaagtgcgaaatatacatagaatgaacaataatccgagctatattatgggcaatattttggctagttttctgcttacaataatacataagaggcctacactaaatacggcggtatataaaactatgcaagacatacccatatattagttttaatcaaattgatttaaaatatttttaaaactattaggcaactttgagtagaataagggtttaacctaaacgtatgtggttttagccatcgatggaaacgttttacaattatcgttacttttttgaagaccttttcgttctatttttaagaagaacattcaaagaaatggaaaaccgttttcttacgaatcctatataccgttgttaattgtttaaaaacacgataaggatattatgttctgaatgtgtcccatcttaccccacagcactatataataaaattttagtttttgtgacagacttatgtcgcattcttaaacggatgatccccaactggttaacaacgtaacaagcttgcaaaagcgaaggtgacattggtaacaacgtacgataactattgtctactaataacaatagacttaacacataacatatttaggaaatggcttcatatggcggactgtcaacttagttgtcaaaattacctttaaatctctataaacgaagctgtttaataaaaaaaactaacaaggctattctacatcaaaccataataatgaaattatcaaacacatttttaaagcgatttcatcaaaccaacgcgccacatgcaagacggtatgcgtcacactgagttttggagtgttctgcatgcgctagacttgaatcagcaggagagttcggaggcttatcaggaagcagttgtccttgttaatgactcgttaagatcaaagtggcatcgaaaacgagtctcgctataaaacgggtttagtttcacagtacctcattccgctttctgttctcattggatatacaccaaactgaaagtacgagttaaatcgaaagagagaggaaattgtaagttttattggaccagacaagactatggcgaatatggcggccgcaaccATGatagttttgcgtttactggttatgggtgcgcttgcgtatgtgagcgtggcaaaaacccggtacgatttatcaggagatataatgcaggggtatgacgctaaagtacgaccgagtgacagctacaataactcggtaaaagtcgtgtttaagcttgtgtttaatcaactactagacgtgagcgaagtcaaccaaaaaatcgaaacgaaactgtgggtttatcacaagtggatggaccctcggttaagctgggtgccagaggattacgaaaacctggagtatatatatctacctactaccaacttgtggctgccggagttggttctgtataacaacgccgatggtgactttgctatttctcaatttacgaaagcaaaagtggattatactggaatggtagaatggaaacctccggcaatttttaaaagcttctgcgagattatggtggcagagtttccatttgacacacaaaactgtacgatgaaaattggcccttggtcgcaaggacaagatttactggacatggtaaattcagactgggaagtcaaagaccacatgtgcgagcccccagatgaaacaatgtacgaggaaagcggagaatggttgattcttaaaactggttgctggaagcattacattaaatacgattgctgtcgagggccgtacgtggacatgacttactacttcatattgcaaagacggcctctatatcttgttatcaacattctctttcctacaatgctgttttcgtatctaacctgcgctgtattctatctgccatcggacgctggcgaaaaaataacactcagtatttcgcttctgctttcactgattgtgttcttgctcgttattgttgaagcgattccctcaaccgctaatggagtcccattgctatgccagtacattctatttactatgatattggtttgcctctctattatgataacagtcggtgtattgaatgtacattaccgtgggcctgcaacacatgtcatgtcggatcggatgaaaaagatattcatggtgtggcttccaaaattcatctatagctcgacaatgaaacgattggatccttacaaggaagaaaaaatgttggctggcagacctcccaaaccttacaaagatatttcagatttatctggtcgGGCCCCCATGGTGAGCAAGGGCGAGGAGCTGTTCACCGGGGTGGTGCCCATCCTGGTCGAGCTGGACGGCGACGTAAACGGCCACAAGTTCAGCGTGTCCGGCGAGGGCGAGGGCGATGCCACCTACGGCAAGCTGACCCTGAAGTTCATCTGCACCACCGGCAAGCTGCCCGTGCCCTGGCCCACCCTCGTGACCACCCTGACCTACGGCGTGCAGTGCTTCAGCCGCTACCCCGACCACATGAAGCAGCACGACTTCTTCAAGTCCGCCATGCCCGAAGGCTACGTCCAGGAGCGCACCATCTTCTTCAAGGACGACGGCAACTACAAGACCCGCGCCGAGGTGAAGTTCGAGGGCGACACCCTGGTGAACCGCATCGAGCTGAAGGGCATCGACTTCAAGGAGGACGGCAACATCCTGGGGCACAAGCTGGAGTACAACTACAACAGCCACAACGTCTATATCATGGCCGACAAGCAGAAGAACGGCATCAAGGTGAACTTCAAGATCCGCCACAACATCGAGGACGGCAGCGTGCAGCTCGCCGACCACTACCAGCAGAACACCCCCATCGGCGACGGCCCCGTGCTGCTGCCCGACAACCACTACCTGAGCACCCAGTCCGCCCTGAGCAAAGACCCCAACGAGAAGCGCGATCACATGGTCCTGCTGGAGTTCGTGACCGCCGCCGGGATCACTCTCGGCATGGACGAGCTGTACAAGCGttccccaaaaccagacgaaccacatttaattggtagcgacgttaaaacagctatggacggagtagattatgtgtcagagtgttataaggaccaacgcgaaggacagcagaaagaagacgaatggaaatacgtcgctatggtgttggaccattttctgttgtatatctttatactggcttgcgtggttggcacagtcggtatatttggcaagcggttgcttgaatttatgtcagaacaagagctttttaaaaacctaggcaacgagtgcattctgaaatgccaaagtgattaAAGCgaattc

*Tbx6-related.b>Cas9*

Tbx6-related.b driver (Christiaen et al. 2009)

Cas9 (Stolfi et al. 2014

ggcgcgcccaacggagtacgcgtgtcaagtttaatggcgtaattaccgaacaactgttgataagtaatgaggaccccgctgcggtaacctttcgaaattgctggttgcaagcggtgttggtgcgataataaggaaaagcggtaagcgttattttttgatccaccacaaagaccaaacctatattgatacactaaactaaatcaaattcgaacctatccattaaagtgcgaaatatacatagaatgaacaataatccgagctatattatgggcaatattttggctagttttctgcttacaataatacataagaggcctacactaaatacggcggtatataaaactatgcaagacatacccatatattagttttaatcaaattgatttaaaatatttttaaaactattaggcaactttgagtagaataagggtttaacctaaacgtatgtggttttagccatcgatggaaacgttttacaattatcgttacttttttgaagaccttttcgttctatttttaagaagaacattcaaagaaatggaaaaccgttttcttacgaatcctatataccgttgttaattgtttaaaaacacgataaggatattatgttctgaatgtgtcccatcttaccccacagcactatataataaaattttagtttttgtgacagacttatgtcgcattcttaaacggatgatccccaactggttaacaacgtaacaagcttgcaaaagcgaaggtgacattggtaacaacgtacgataactattgtctactaataacaatagacttaacacataacatatttaggaaatggcttcatatggcggactgtcaacttagttgtcaaaattacctttaaatctctataaacgaagctgtttaataaaaaaaactaacaaggctattctacatcaaaccataataatgaaattatcaaacacatttttaaagcgatttcatcaaaccaacgcgccacatgcaagacggtatgcgtcacactgagttttggagtgttctgcatgcgctagacttgaatcagcaggagagttcggaggcttatcaggaagcagttgtccttgttaatgactcgttaagatcaaagtggcatcgaaaacgagtctcgctataaaacgggtttagtttcacagtacctcattccgctttctgttctcattggatatacaccaaactgaaagtacgagttaaatcgaaagagagaggaaattgtaagttttattggaccagacaagactatggcgaatatggcggccgcaaccATGGCTAGCCCCAAAAAGAAGAGGAAAGTGGACAAGAAGTATTCTATCGGACTGGACATCGGGACTAATAGCGTCGGGTGGGCCGTGATCACTGACGAGTACAAGGTGCCCTCTAAGAAGTTCAAGGTGCTCGGGAACACCGACCGGCATTCCATCAAGAAAAATCTGATCGGAGCTCTCCTCTTTGATTCAGGGGAGACCGCTGAAGCAACCCGCCTCAAGCGGACTGCTAGACGGCGGTACACCAGGAGGAAGAACCGGATTTGTTACCTTCAAGAGATATTCTCCAACGAAATGGCAAAGGTCGACGACAGCTTCTTCCATAGGCTGGAAGAATCATTCCTCGTGGAAGAGGATAAGAAGCATGAACGGCATCCCATCTTCGGTAATATCGTCGACGAGGTGGCCTATCACGAGAAATACCCAACCATCTACCATCTTCGCAAAAAGCTGGTGGACTCAACCGACAAGGCAGACCTCCGGCTTATCTACCTGGCCCTGGCCCACATGATCAAGTTCAGAGGCCACTTCCTGATCGAGGGCGACCTCAATCCTGACAATAGCGATGTGGATAAACTGTTCATCCAGCTGGTGCAGACTTACAACCAGCTCTTTGAAGAGAACCCCATCAATGCAAGCGGAGTCGATGCCAAGGCCATTCTGTCAGCCCGGCTGTCAAAGAGCCGCAGACTTGAGAATCTTATCGCTCAGCTGCCGGGTGAAAAGAAAAATGGACTGTTCGGGAACCTGATTGCTCTTTCACTTGGGCTGACTCCCAATTTCAAGTCTAATTTCGACCTGGCAGAGGATGCCAAGCTGCAACTGTCCAAGGACACCTATGATGACGATCTCGACAACCTCCTGGCCCAGATCGGTGACCAATACGCCGACCTTTTCCTTGCTGCTAAGAATCTTTCTGACGCCATCCTGCTGTCTGACATTCTCCGCGTGAACACTGAAATCACCAAGGCCCCTCTTTCAGCTTCAATGATTAAGCGGTATGATGAGCACCACCAGGACCTGACCCTGCTTAAGGCACTCGTCCGGCAGCAGCTTCCGGAGAAGTACAAGGAAATCTTCTTTGACCAGTCAAAGAATGGATACGCCGGCTACATCGACGGAGGTGCCTCCCAAGAGGAATTTTATAAGTTTATCAAACCTATCCTTGAGAAGATGGACGGCACCGAAGAGCTCCTCGTGAAACTGAATCGGGAGGATCTGCTGCGGAAGCAGCGCACTTTCGACAATGGGAGCATTCCCCACCAGATCCATCTTGGGGAGCTTCACGCCATCCTTCGGCGCCAAGAGGACTTCTACCCCTTTCTTAAGGACAACAGGGAGAAGATTGAGAAAATTCTCACTTTCCGCATCCCCTACTACGTGGGACCCCTCGCCAGAGGAAATAGCCGGTTTGCTTGGATGACCAGAAAGTCAGAAGAAACTATCACTCCCTGGAACTTCGAAGAGGTGGTGGACAAGGGAGCCAGCGCTCAGTCATTCATCGAACGGATGACTAACTTCGATAAGAACCTCCCCAATGAGAAGGTCCTGCCGAAACATTCCCTGCTCTACGAGTACTTTACCGTGTACAACGAGCTGACCAAGGTGAAATATGTCACCGAAGGGATGAGGAAGCCCGCATTCCTGTCAGGCGAACAAAAGAAGGCAATTGTGGACCTTCTGTTCAAGACCAATAGAAAGGTGACCGTGAAGCAGCTGAAGGAGGACTATTTCAAGAAAATTGAATGCTTCGACTCTGTGGAGATTAGCGGGGTCGAAGATCGGTTCAACGCAAGCCTGGGTACCTACCATGATCTGCTTAAGATCATCAAGGACAAGGATTTTCTGGACAATGAGGAGAACGAGGACATCCTTGAGGACATTGTCCTGACTCTCACTCTGTTCGAGGACCGGGAAATGATCGAGGAGAGGCTTAAGACCTACGCCCATCTGTTCGACGATAAAGTGATGAAGCAACTTAAACGGAGAAGATATACCGGATGGGGACGCCTTAGCCGCAAACTCATCAACGGAATCCGGGACAAACAGAGCGGAAAGACCATTCTTGATTTCCTTAAGAGCGACGGATTCGCTAATCGCAACTTCATGCAACTTATCCATGATGATTCCCTGACCTTTAAGGAGGACATCCAGAAGGCCCAAGTGTCTGGACAAGGTGACTCACTGCACGAGCATATCGCAAATCTGGCTGGTTCACCCGCTATTAAGAAGGGTATTCTCCAGACCGTGAAAGTCGTGGACGAGCTGGTCAAGGTGATGGGTCGCCATAAACCAGAGAACATTGTCATCGAGATGGCCAGGGAAAACCAGACTACCCAGAAGGGACAGAAGAACAGCAGGGAGCGGATGAAAAGAATTGAGGAAGGGATTAAGGAGCTCGGGTCACAGATCCTTAAAGAGCACCCGGTGGAAAACACCCAGCTTCAGAATGAGAAGCTCTATCTGTACTACCTTCAAAATGGACGCGATATGTATGTGGACCAAGAGCTTGATATCAACAGGCTCTCAGACTACGACGTGGACCACATCGTCCCTCAGAGCTTCCTCAAAGACGACTCAATTGACAATAAGGTGCTGACTCGCTCAGACAAGAACCGGGGAAAGTCAGATAACGTGCCCTCAGAGGAAGTCGTGAAAAAGATGAAGAACTATTGGCGCCAGCTTCTGAACGCAAAGCTGATCACTCAGCGGAAGTTCGACAATCTCACTAAGGCTGAGAGGGGCGGACTGAGCGAACTGGACAAAGCAGGATTCATTAAACGGCAACTTGTGGAGACTCGGCAGATTACTAAACATGTCGCCCAAATCCTTGACTCACGCATGAATACCAAGTACGACGAAAACGACAAACTTATCCGCGAGGTGAAGGTGATTACCCTGAAGTCCAAGCTGGTCAGCGATTTCAGAAAGGACTTTCAATTCTACAAAGTGCGGGAGATCAATAACTATCATCATGCTCATGACGCATATCTGAATGCCGTGGTGGGAACCGCCCTGATCAAGAAGTACCCAAAGCTGGAAAGCGAGTTCGTGTACGGAGACTACAAGGTCTACGACGTGCGCAAGATGATTGCCAAATCTGAGCAGGAGATCGGAAAGGCCACCGCAAAGTACTTCTTCTACAGCAACATCATGAATTTCTTCAAGACCGAAATCACCCTTGCAAACGGTGAGATCCGGAAGAGGCCGCTCATCGAGACTAATGGGGAGACTGGCGAAATCGTGTGGGACAAGGGCAGAGATTTCGCTACCGTGCGCAAAGTGCTTTCTATGCCTCAAGTGAACATCGTGAAGAAAACCGAGGTGCAAACCGGAGGCTTTTCTAAGGAATCAATCCTCCCCAAGCGCAACTCCGACAAGCTCATTGCAAGGAAGAAGGATTGGGACCCTAAGAAGTACGGCGGATTCGATTCACCAACTGTGGCTTATTCTGTCCTGGTCGTGGCTAAGGTGGAAAAAGGAAAGTCTAAGAAGCTCAAGAGCGTGAAGGAACTGCTGGGTATCACCATTATGGAGCGCAGCTCCTTCGAGAAGAACCCAATTGACTTTCTCGAAGCCAAAGGTTACAAGGAAGTCAAGAAGGACCTTATCATCAAGCTCCCAAAGTATAGCCTGTTCGAACTGGAGAATGGGCGGAAGCGGATGCTCGCCTCCGCTGGCGAACTTCAGAAGGGTAATGAGCTGGCTCTCCCCTCCAAGTACGTGAATTTCCTCTACCTTGCAAGCCATTACGAGAAGCTGAAGGGGAGCCCCGAGGACAACGAGCAAAAGCAACTGTTTGTGGAGCAGCATAAGCATTATCTGGACGAGATCATTGAGCAGATTTCCGAGTTTTCTAAACGCGTCATTCTCGCTGATGCCAACCTCGATAAAGTCCTTAGCGCATACAATAAGCACAGAGACAAACCAATTCGGGAGCAGGCTGAGAATATCATCCACCTGTTCACCCTCACCAATCTTGGTGCCCCTGCCGCATTCAAGTACTTCGACACCACCATCGACCGGAAACGCTATACCTCCACCAAAGAAGTGCTGGACGCCACCCTCATCCACCAGAGCATCACCGGACTTTACGAAACTCGGATTGACCTCTCACAGCTCGGAGGGGATGAGGGAGCTCCCAAGAAAAAGCGCAAGGTAGGTTAATGAgaattc

*Tbx6-related.b>CD4::mCherry*

Tbx6-related.b driver (Christiaen et al. 2009)

CD4::mCherry, human (Gline et al. 2015)

ggcgcgcccaacggagtacgcgtgtcaagtttaatggcgtaattaccgaacaactgttgataagtaatgaggaccccgctgcggtaacctttcgaaattgctggttgcaagcggtgttggtgcgataataaggaaaagcggtaagcgttattttttgatccaccacaaagaccaaacctatattgatacactaaactaaatcaaattcgaacctatccattaaagtgcgaaatatacatagaatgaacaataatccgagctatattatgggcaatattttggctagttttctgcttacaataatacataagaggcctacactaaatacggcggtatataaaactatgcaagacatacccatatattagttttaatcaaattgatttaaaatatttttaaaactattaggcaactttgagtagaataagggtttaacctaaacgtatgtggttttagccatcgatggaaacgttttacaattatcgttacttttttgaagaccttttcgttctatttttaagaagaacattcaaagaaatggaaaaccgttttcttacgaatcctatataccgttgttaattgtttaaaaacacgataaggatattatgttctgaatgtgtcccatcttaccccacagcactatataataaaattttagtttttgtgacagacttatgtcgcattcttaaacggatgatccccaactggttaacaacgtaacaagcttgcaaaagcgaaggtgacattggtaacaacgtacgataactattgtctactaataacaatagacttaacacataacatatttaggaaatggcttcatatggcggactgtcaacttagttgtcaaaattacctttaaatctctataaacgaagctgtttaataaaaaaaactaacaaggctattctacatcaaaccataataatgaaattatcaaacacatttttaaagcgatttcatcaaaccaacgcgccacatgcaagacggtatgcgtcacactgagttttggagtgttctgcatgcgctagacttgaatcagcaggagagttcggaggcttatcaggaagcagttgtccttgttaatgactcgttaagatcaaagtggcatcgaaaacgagtctcgctataaaacgggtttagtttcacagtacctcattccgctttctgttctcattggatatacaccaaactgaaagtacgagttaaatcgaaagagagaggaaattgtaagttttattggaccagacaagactatggcgaatatggcggccgcaaccATGAACCGGGGAGTCCCTTTTAGGCACTTGCTTCTGGTGCTGCAACTGGCGCTCCTCCCAGCAGCCACTCAGGGAAAGAAAGTGGTGCTGGGCAAAAAAGGGGATACAGTGGAACTGACCTGTACAGCTTCCCAGAAGAAGAGCATACAATTCCACTGGAAAAACTCCAACCAGATAAAGATTCTGGGAAATCAGGGCTCCTTCTTAACTAAAGGTCCATCCAAGCTGAATGATCGCGCTGACTCAAGAAGAAGCCTTTGGGACCAAGGAAACTTCCCCCTGATCATCAAGAATCTTAAGATAGAAGACTCAGATACTTACATCTGTGAAGTGGAGGACCAGAAGGAGGAGGTGCAATTGCTAGTGTTCGGATTGACTGCCAACTCTGACACCCACCTGCTTCAGGGGCAGAGCCTGACCCTGACCTTGGAGAGCCCCCCTGGTAGTAGCCCCTCAGTGCAATGTAGGAGTCCAAGGGGTAAAAACATACAGGGGGGGAAGACCCTCTCCGTGTCTCAGCTGGAGCTCCAGGATAGTGGCACCTGGACATGCACTGTCTTGCAGAACCAGAAGAAGGTGGAGTTCAAAATAGACATCGTGGTGCTAGCTTTCCAGAAGGCCTCCAGCATAGTCTATAAGAAAGAGGGGGAACAGGTGGAGTTCTCCTTCCCACTCGCCTTTACAGTTGAAAAGCTGACGGGCAGTGGCGAGCTGTGGTGGCAGGCGGAGAGGGCTTCCTCCTCCAAGTCTTGGATCACCTTTGACCTGAAGAACAAGGAAGTGTCTGTAAAACGGGTTACCCAGGACCCTAAGCTCCAGATGGGCAAGAAGCTCCCGCTCCACCTCACCCTGCCCCAGGCCTTGCCTCAGTATGCTGGCTCTGGAAACCTCACCCTGGCCCTTGAAGCGAAAACAGGAAAGTTGCATCAGGAAGTGAACCTGGTGGTGATGAGAGCCACTCAGCTCCAGAAAAATTTGACCTGTGAGGTGTGGGGACCCACCTCCCCTAAGCTGATGCTGAGCTTGAAACTGGAGAACAAGGAGGCAAAGGTCTCGAAGCGGGAGAAGGCGGTGTGGGTGCTGAACCCTGAGGCGGGGATGTGGCAGTGTCTGCTGAGTGACTCGGGACAGGTCCTGCTGGAATCCAACATCAAGGTTCTGCCCACATGGTCCACCCCGGTGCAGCCAATGGCCCTGATTGTGCTGGGGGGCGTCGCCGGCCTCCTGCTTTTCATTGGGCTAGGCATCTTCTTCTGTGTCAGGACTAGTGTGAGCAAGGGCGAGGAGGATAACATGGCCATCATCAAGGAGTTCATGCGCTTCAAGGTGCACATGGAGGGCTCCGTGAACGGCCACGAGTTCGAGATCGAGGGCGAGGGCGAGGGCCGCCCCTACGAGGGCACCCAGACCGCCAAGCTGAAGGTGACCAAGGGTGGCCCCCTGCCCTTCGCCTGGGACATCCTGTCCCCTCAGTTCATGTACGGCTCCAAGGCCTACGTGAAGCACCCCGCCGACATCCCCGACTACTTGAAGCTGTCCTTCCCCGAGGGCTTCAAGTGGGAGCGCGTGATGAACTTCGAGGACGGCGGCGTGGTGACCGTGACCCAGGACTCCTCCCTGCAGGACGGCGAGTTCATCTACAAGGTGAAGCTGCGCGGCACCAACTTCCCCTCCGACGGCCCCGTAATGCAGAAGAAGACCATGGGCTGGGAGGCCTCCTCCGAGCGGATGTACCCCGAGGACGGCGCCCTGAAGGGCGAGATCAAGCAGAGGCTGAAGCTGAAGGACGGCGGCCACTACGACGCTGAGGTCAAGACCACCTACAAGGCCAAGAAGCCCGTGCAGCTGCCCGGCGCCTACAACGTCAACATCAAGTTGGACATCACCTCCCACAACGAGGACTACACCATCGTGGAACAGTACGAACGCGCCGAGGGCCGCCACTCCACCGGCGGCATGGACGAGCTGTACAAGTAAgaattc

*Islet -7216/-3950 + bpFOG>tagRFP*

Islet -7216/-3950 (Stolfi et al. 2010)

bpFOG (Rothbächer et al. 2007)

TagRFP (Merzlyak et al. 2007)

ggcgcgccaagtataccacgcgagttagttcggccaggaagcgcgctgtttattatcaagttattatgacgtcatataaaatacgtcacatttgtagcacctggttatctgccttcccacggttaaaacagatgcgacctcggttatcagtggtttgacatggagatcgtaaatcaagcgacgaatatctgtgacacgtcacgtgacgagattgctgacataccaggacacttaacggtatctggttaggggaagtacctgtagcgtcatacacataacctgcataatgctttatagcttcgaatacgatagcgaagatcaaattaattaaaacaacgaagagaaggattcgaagcaaaacgacagtcgttaaaacacactacgggtaaaaattacttggggaaaagcgtggctgttacatccaaatcaaaatgtaagccgtcaatggtttgtgaacaaacggtaactcttgtcacctctaagctaaactcattttttgcagagtttaaaaataccgagtagcaactgccgtctggaacgaataaatgaatctaacgacgcagacgttatgacaggtgtgttcatacaccatgtttcccttttacgggaaagcgtataaaatatatatttacattcttgcctaaagtataaacgaacttttattgaatggtcaaaatgggtcatcttaaattccattataacttttccatttcaaacatttttccgcagcatcgtcataaaacagggattccaatagaaacatcataaaacaccaaaccatgttgtttatggatcaccgtgtttgttgtattaacaagagttataactacaacatgacgtgactagattgacacaacaactatggttggcagtcaaaacacacgaccccaccccataagatggagccacgccaccaaatcaacacagcagaagctccaggtggtttggtaggttctgttatcacaacactacaggccatgagatctatgatctataccgtctgttatatttagtggatttgtgattttaaccgaacaccttcactagtttaaaccaaccgacaactcaccgcgaacaatcagcgaatatttcattaaaccgacaaaaatttccagttaatttaaaatatagtaagttggggtaagatgggacatgtttttattctcatctcgtcccatttggtagtcaacaaagaatatttacagaattataaattgaaatatcttcacgaagagaaaaatattgttatttgttaaaaacacgatcaggaaatatgggatattttgtgctaaccgtattcatatttgcaacccactgtgagttgcttcttataaggtttaacagccacggttttaactaaattttatttattttaaccttagcgtgttataacgactgttatttttctgttaattcccatttcttcgccatgttagttaatatgacttttgttattgtaattatataaaaatatattttttaaaataagtctaaaagtaaatgtctaagttgtatttacaaatttccgcaccagaataggcagcaggcaaccgcggatcgaaacaagtccctgatttatggttttgttgcaactaattatggctgtacagcacacaggtcccaggattgccgtaaaacttaatttgctggaatatgttgaacaacagttgttacattctgttgttttattaagttcagcggccacgcttttttaagccttttattccagtattttcgctcgtagcgtgttgataacgactgttgttttgctgctaattctcttcgttgttttaattaaacttcgctatcgtattcaaaatatgttcatttagttggtaggttatgtgtgtaagtgacatcataacaacttgtatgaggcaaacaaggggttatgacgtaataatagaattattgatgtttcacagggtgaggtgtgacgtaataatagttctgacagttaaagtgattttgaataaaaacaaaaaacagaatgtttaagtcgcgaaaatcgcgtgggcagcgttaaccagagcatcgcatgtgacaagcgaacgaaatcccagcagatctacgacgttggtccgctgagagttcgagcgaattcgggatgtaacgtcgatgaaagacgaaaagcggtccttgcctcgacaagaaaggacgaaccgcgcgatagtagattttatggcgtatggtgttctataacacagctgtgggttgaagtggtttcgaatatttaagcaaggtacgacttgaccggttacgtcactttgaccacgtggtcagctctcatcaatatttattttaaatttgattaaacagcgacctcaggtgagcggcctaattatgcgcctctatgacgtcatagtttaaacttcttttaatagcggaatcgaagataaataggttgcgttaccctgggggtcgagatccaaatttcgttacttcgattgtttaattgattcgacatcgagttaaagcgcgagatttgattacatttccattgttcaatccataattcgctttgtgacgtaacacaaacttaacaaacacgtggtagcgcgcaagagctaattttttaaacgtttcttgattttttttcgattttttttattaaaaaataaaataaattaatattttgttgaaatacgtcgaaaccgcgaaaacgcgatcgacgcaattcaatttgtaaatgaaatatcgaccagtggcgccgatagcggataatcctcacaattcttgtgataaattagtttcttactcggataatctgtgtgccggtttgattgtcgcacccaaggtgattggggggggggggatattacgtcattatgacaaataatttcgattaaaattggtaaaaatataaattaggggagtttttagttttgttttaaaagattttatttaaaaattaaaaaaaaaatcgaaaatttttgaatttgatcgattagctgtaattacgtcatcaatccgctcattatgacgtcatacgtcgccatattgcgtcataatcgggcgtcagaatgtcgtcacaaaccatattgtgttgaattaattggaattatgaaacgccgctcggtcgggggatttgtgacgtcacaaacgtgaatgttacgacacaaacggcgaactcgggcgaacttttagctttgcggataataagggtaagtgacgcaacaatgacgcaacaatgttgtcccttgtgacgtaacagagtgtattgtaaccgttgaattgagtctcgagcagctgaagcttgcatgcctgcaggtcgactctagaggatccggcaaAGCTTCGTGTATTGTACCGGCCCATTGTCAATCATGCAAACTTGATATTATATTGACAAGAGAAGAAGGCAGTTTAAATTAAAACTCTAAAGTAGAGAGACATTAATCTCAGCTGACAAGGCAGGTGGTCACAGTAAGTTCATTTAAATAGTTGGCCAACAATAGCCTTTCCAAGAAAGTATTTTTGTTCCAGGTCTATACAAAAATAACACACATAgcggccgcaaccATGGTGTCTAAGGGCGAAGAGCTGATTAAGGAGAACATGCACATGAAGCTGTACATGGAGGGCACCGTGAACAACCACCACTTCAAGTGCACATCCGAGGGCGAAGGCAAGCCCTACGAGGGCACCCAGACCATGAGAATCAAGGTGGTCGAGGGCGGCCCTCTCCCCTTCGCCTTCGACATCCTGGCTACCAGCTTCATGTACGGCAGCAGAACCTTCATCAACCACACCCAGGGCATCCCCGACTTCTTTAAGCAGTCCTTCCCTGAGGGCTTCACATGGGAGAGAGTCACCACATACGAAGACGGGGGCGTGCTGACCGCTACCCAGGACACCAGCCTCCAGGACGGCTGCCTCATCTACAACGTCAAGATCAGAGGGGTGAACTTCCCATCCAACGGCCCTGTGATGCAGAAGAAAACACTCGGCTGGGAGGCCAACACCGAGATGCTGTACCCCGCTGACGGCGGCCTGGAAGGCAGAAGCGACATGGCCCTGAAGCTCGTGGGCGGGGGCCACCTGATCTGCAACTTCAAGACCACATACAGATCCAAGAAACCCGCTAAGAACCTCAAGATGCCCGGCGTCTACTATGTGGACCACAGACTGGAAAGAATCAAGGAGGCCGACAAAGAGACCTACGTCGAGCAGCACGAGGTGGCTGTGGCCAGATACTGCGACCTCCCTAGCAAACTGGGGCACAACTTAATTAAgaattc

*Eef1a>H2B::CFP*

Eef1a -1955/-1 (Sasakura et al. 2010)

H2B::CFP (Stolfi et al. 2010)

ggcgcgccgtgacgggaaaacgatagtcgttataacacgagtattcgtacacctcgtgcgagctaacgagctaccatatatgttgtgggcgaataaaggttttataaatataacattggttttataaataaaacaacgccattttaaagtcggttacataattctgtaactagttcaaattgaacggtaaacgtaaataaaaaccttgaccgtcttacccaattatataaaaacactttgaacgctttttaagatggaagggtatggccatgcctagataattctgtggaccatctcaccccaacctattacagaacggtcgtaataatgaaaatgggtaccatttttaggcatatagactgattcctcctttctagaaacgtaagcagtatacacagaaaaaatgaagtgtgattctgtgcaattaaaccgttctaaattcatagccgactgaatttctaattaagtgaatgtctgacctagatttattgttaagtttagcaccaaatctgagccagcgataagcagtctaattaaattggctgctggcgataaaataggtcatcctgaaaaatcgtttgcgcctttatttaaaatatagtagagtggggaaagacgggacatcttatcgttctattttctcgtcccatttcgtagtaaacaaagaacattcaaaaaatataaaaccataacttcaaaacttcaatagaccgttgtcaactgtttaaaacacaataagagaatttggatattatgtgctaaaggtgtcccatctccccccaccctactatatctgtttatagttctgtggggtaagatgagataccgttaacacctaaacatttttactttaaacaatcaaccacgttttttatagtcgtaatggacatgtggttacataattctgaaaatattttttgcccccgaccaaaagacgcgaagagtaaaaacatgtctcagcttatattccccacataaatatatttttgtactgtttggtgaatttataaacttatattaccatgcatatacgttatgttactggtattttctcagtaggcaaattcatttgtccacgttttataggttttcaatatttatgatttttaaaatgctaaaaatgtgggaggggggttgaaagtacaatacaaacacacaaaacaactcaaactaaagatttatagttatgctaattcacctacacaatataacaagatgtgtaatgcaaccatgtgtttatgatgagcgctaacatattttgtaaccactcaaattccccgccacacgaggataatgaataggtgactctgtagtctgtacatcttagactgaaataaagattataaatctacgaaataaaataatttctgctcactgattatacttctgttttatagattagaaaccgtttctaataaatgacctaattcgctatacacacacgctgtgcgcgagataatcattctcgcaccccgtttattgtgttaaaattgccgcctagattcacaaagcgtgacggctagagccagcaacgtgtcgccttcaattacgcaacatccgggttgcgcaattctggatataaaagaactaacaaagatgacgtagctacctttttcagttcagacttacgaaagactcacgtgtcggcggtctacttgtccttttcgagctgtggcaatttggtgagtggttctatcttatatctgagtacatctctaaggaattatagtttgattagttaagtttttattgttaggaaagatgaaatcattaggttttacttagtttaagtatgttagtactggttaggcgtttgaattattgaaaaactcagttcgttaactgtagtagttctggtagcttagcaagtataccctgtatacgccttttggctttttaacaataacttaaacttattttacagcaaatttctgtgcattcggttaaccccaaccttccaaagcggccgcAACCATGGTTGCATCCAAAGGTCCCAAGAAAGCTATGAAGAGCACACCCCCAGTAGAGAAGAAGGAACGCAAGAATAAGCGCCATCGACGACGAAAGGAAAGCTTCGGCATCTACATTTACAAAGTGCTGAAGCAAGTCCACCCTGACACTGGTATCTCCAGCAAGGCAATGAACATCATGAATTCATTTGTTCACGACCTATTCGAAAGAGTTGCCGGAGAATCTTCTCGTCTCTGTAGCTACAACAAGCGCTCCACTATCAGCAGTCGGGAAGTGCAAACCGCAGTTCGCCTTTTGCTTCCAGGGGAGCTTGCAAAACATGCTGTATCGGAGGGTACAAAGGCCGTTACAAAGTACACCAGCTCAAAAACTAGTTCCAAAGGTGAAGAACTTTTCACTGGAGTGGTGCCTATCTTGGTTGAGCTTGACGGTGATGTGAACGGTCACAAATTCTCTGTAAGTGGTGAAGGAGAGGGAGACGCTACCTACGGCAAGTTAACGCTGAAATTTATATGCACTACGGGAAAGCTGCCTGTACCGTGGCCTACACTGGTTACCACTCTGACTTGGGGAGTACAATGCTTCGCCCGCTATCCGGACCACATGAAACGCCATGACTTCTTTAAATCAGCTATGCCAGAAGGATACGTGCAAGAACGAACCATTTTCTTCAAAGATGATGGTAATTATAAAACAAGGGCGGAAGTTAAATTCGAAGGAGACACGCTCGTAAACAGAATCGAACTTAAGGGTATCGACTTCAAGGAGGATGGCAACATTCTTGGACACAAACTGGAGTACAACGCCATTTCCGATAATGTTTACATTACTGCTGATAAACAGAAGAACGGAATTAAGGCGAATTTTAAAATCAGACATAACATTGAAGATGGTGGAGTGCAATTGGCTGATCACTACCAGCAAAATACTCCTATCGGAGACGGCCCTGTGTTGCTTCCTGACAACCACTACTTAAGTACGCAATCAGCTTTATCCAAAGATCCCAATGAAAAGCGAGATCACATGGTGCTGCTCGAATTTGTTACTGCTGCTGGTATTACACACGGAATGGACGAGCTGTACAAGTAAgaattc

**New constructs:**

*Eef1a>Nova(MLN)*

Eef1a -1955/-1 (Sasakura et al. 2010)

Nova(MLN)

ggcgcgccgtgacgggaaaacgatagtcgttataacacgagtattcgtacacctcgtgcgagctaacgagctaccatatatgttgtgggcgaataaaggttttataaatataacattggttttataaataaaacaacgccattttaaagtcggttacataattctgtaactagttcaaattgaacggtaaacgtaaataaaaaccttgaccgtcttacccaattatataaaaacactttgaacgctttttaagatggaagggtatggccatgcctagataattctgtggaccatctcaccccaacctattacagaacggtcgtaataatgaaaatgggtaccatttttaggcatatagactgattcctcctttctagaaacgtaagcagtatacacagaaaaaatgaagtgtgattctgtgcaattaaaccgttctaaattcatagccgactgaatttctaattaagtgaatgtctgacctagatttattgttaagtttagcaccaaatctgagccagcgataagcagtctaattaaattggctgctggcgataaaataggtcatcctgaaaaatcgtttgcgcctttatttaaaatatagtagagtggggaaagacgggacatcttatcgttctattttctcgtcccatttcgtagtaaacaaagaacattcaaaaaatataaaaccataacttcaaaacttcaatagaccgttgtcaactgtttaaaacacaataagagaatttggatattatgtgctaaaggtgtcccatctccccccaccctactatatctgtttatagttctgtggggtaagatgagataccgttaacacctaaacatttttactttaaacaatcaaccacgttttttatagtcgtaatggacatgtggttacataattctgaaaatattttttgcccccgaccaaaagacgcgaagagtaaaaacatgtctcagcttatattccccacataaatatatttttgtactgtttggtgaatttataaacttatattaccatgcatatacgttatgttactggtattttctcagtaggcaaattcatttgtccacgttttataggttttcaatatttatgatttttaaaatgctaaaaatgtgggaggggggttgaaagtacaatacaaacacacaaaacaactcaaactaaagatttatagttatgctaattcacctacacaatataacaagatgtgtaatgcaaccatgtgtttatgatgagcgctaacatattttgtaaccactcaaattccccgccacacgaggataatgaataggtgactctgtagtctgtacatcttagactgaaataaagattataaatctacgaaataaaataatttctgctcactgattatacttctgttttatagattagaaaccgtttctaataaatgacctaattcgctatacacacacgctgtgcgcgagataatcattctcgcaccccgtttattgtgttaaaattgccgcctagattcacaaagcgtgacggctagagccagcaacgtgtcgccttcaattacgcaacatccgggttgcgcaattctggatataaaagaactaacaaagatgacgtagctacctttttcagttcagacttacgaaagactcacgtgtcggcggtctacttgtccttttcgagctgtggcaatttggtgagtggttctatcttatatctgagtacatctctaaggaattatagtttgattagttaagtttttattgttaggaaagatgaaatcattaggttttacttagtttaagtatgttagtactggttaggcgtttgaattattgaaaaactcagttcgttaactgtagtagttctggtagcttagcaagtataccctgtatacgccttttggctttttaacaataacttaaacttattttacagcaaatttctgtgcattcggttaaccccaaccttccaaagcggccgcAACCATGCTAAATGCAATGGAGTATGAATGCCAGTACAATGCTGGCTACAGCATTGTGTCTAACGGTAACGAATACGGTCTCATACAGGCCTACACGGCACACGATTACCCCCTTGAAAACGGAGTGACGTTTTCAGCACCTCCGCCGGGCCAGCTCATTCTTAAAGTTCTAATACCGGGGTACGCTGCGGGGGCGGTGATCGGGAAAGGCGGTCAGATTATTGTACAACTTCAGAAAGATTCAGGGGCCATTATTAAGCTGTCAAAAGCGAAGGACTTTTACCCCGGAACCCAAGACCGAGTCGTTTTGATCCAAGGAACCGCCGAAGGCTTGATGAAGGTGCAAAATACCATTATAGAGAAGGTGTACGAGTTCCCTGTGCCCAAAGATTTAGCTGCGATCATCGGAGACCGACCGAAACAGGTGAAAATCATCGTACCCAACACAACTGCGGGACTGGTAATAGGAAAGGCCGGCGCAACGATAAAGACCATTATGGAAGAGAGTGGATCGAAGGTTCAACTCTCGCAAAAGCCAGACGGGGTAAACGTCCAAGAACGAGTCATCACAATCAAAGGAGAGAAGCACCAACTCATGACAGCATCTAATATTATTATTGATAAAATTAAAGACGACCCTCAAAGCGCCAGTTGCCCTCACATAAGTTACTCTGGCATCGCTGGCCCGATCGCTAACGCGAATCCCACCGGATCGCCCTACGCTGCTGGCTCGGCTGCATTAGTTGACGCTTCGCACCCATCCGTGGCCGCTATGTTGGGACATTATGTTATCCCAGGCCAACAGGTGCTGCAGACAGCAATGCCACTCTCCCATCACCCGCACCAGTCCGCGTTGTCCAGCGGCTCAGTGACACCGGCGCCTGAACTGACGACCATAAATCACGCCATGACAACGTTAGCGAACTATGGCTACACCTTAGGAGGCGTGAACTATGGTACCTTGGGTGTAATGCCTAGTGTACATCCAAGTGTACACCCTGGCATCGCTACCTCGGTCGGGATGATCTCTGCAGGCTCCCTAGCAGGAAGTCCAATCCCTTCAGCTACCCCCTTGCTCTCTGCCACTGCTCTACCGACGGAATCCAGTATTCCGACGGCTGTTCCCACTGCCCAAGCCATTTCAATGCAGAGCAATTACCTTGCAAACTTGGCTAATGCTGGTTACCTGACTACCGGTCACCCACAGTTGCTTGGAGCGACTATCCTAAGCATCGAAAAGTCAAGCGACGGACAAAAAGAAACAATTGAACTGGCAATTCCCGAAAACCTGATCGGAGCAGTCCTCGGAAAAGCGGGAAGGACACTGGTTGAGTATCAGGATGTATCAGGGGCGAAAATTCAAATTTCTAAAAAGGGTGATTACGTCGCCGGGACCAGGAACAGGAGGGTTACGATTACGGGGAAGCCCCCATGCCCACAGACTGCGCAGTTTCTTATTACGCAACGTGTCGCCTCTGCGCAAAACGCAAGGGCACAGCAGGCTAAGTTACTGTAGgaattc

*Eef1a>Nova(MMM)*

Eef1a -1955/-1 (Sasakura et al. 2010)

Nova(MMM)

ggcgcgccgtgacgggaaaacgatagtcgttataacacgagtattcgtacacctcgtgcgagctaacgagctaccatatatgttgtgggcgaataaaggttttataaatataacattggttttataaataaaacaacgccattttaaagtcggttacataattctgtaactagttcaaattgaacggtaaacgtaaataaaaaccttgaccgtcttacccaattatataaaaacactttgaacgctttttaagatggaagggtatggccatgcctagataattctgtggaccatctcaccccaacctattacagaacggtcgtaataatgaaaatgggtaccatttttaggcatatagactgattcctcctttctagaaacgtaagcagtatacacagaaaaaatgaagtgtgattctgtgcaattaaaccgttctaaattcatagccgactgaatttctaattaagtgaatgtctgacctagatttattgttaagtttagcaccaaatctgagccagcgataagcagtctaattaaattggctgctggcgataaaataggtcatcctgaaaaatcgtttgcgcctttatttaaaatatagtagagtggggaaagacgggacatcttatcgttctattttctcgtcccatttcgtagtaaacaaagaacattcaaaaaatataaaaccataacttcaaaacttcaatagaccgttgtcaactgtttaaaacacaataagagaatttggatattatgtgctaaaggtgtcccatctccccccaccctactatatctgtttatagttctgtggggtaagatgagataccgttaacacctaaacatttttactttaaacaatcaaccacgttttttatagtcgtaatggacatgtggttacataattctgaaaatattttttgcccccgaccaaaagacgcgaagagtaaaaacatgtctcagcttatattccccacataaatatatttttgtactgtttggtgaatttataaacttatattaccatgcatatacgttatgttactggtattttctcagtaggcaaattcatttgtccacgttttataggttttcaatatttatgatttttaaaatgctaaaaatgtgggaggggggttgaaagtacaatacaaacacacaaaacaactcaaactaaagatttatagttatgctaattcacctacacaatataacaagatgtgtaatgcaaccatgtgtttatgatgagcgctaacatattttgtaaccactcaaattccccgccacacgaggataatgaataggtgactctgtagtctgtacatcttagactgaaataaagattataaatctacgaaataaaataatttctgctcactgattatacttctgttttatagattagaaaccgtttctaataaatgacctaattcgctatacacacacgctgtgcgcgagataatcattctcgcaccccgtttattgtgttaaaattgccgcctagattcacaaagcgtgacggctagagccagcaacgtgtcgccttcaattacgcaacatccgggttgcgcaattctggatataaaagaactaacaaagatgacgtagctacctttttcagttcagacttacgaaagactcacgtgtcggcggtctacttgtccttttcgagctgtggcaatttggtgagtggttctatcttatatctgagtacatctctaaggaattatagtttgattagttaagtttttattgttaggaaagatgaaatcattaggttttacttagtttaagtatgttagtactggttaggcgtttgaattattgaaaaactcagttcgttaactgtagtagttctggtagcttagcaagtataccctgtatacgccttttggctttttaacaataacttaaacttattttacagcaaatttctgtgcattcggttaaccccaaccttccaaagcggccgcAACCatgatgatgacggccgtagtacctatgccgaacgggacatatcttatcgagtcgcgaaaaagaccgctggaagaaccgattgaactggtcgatttcaagagagaacgctccgaaatggaagATTACCCCCTTGAAAACGGAGTGACGTTTTCAGCACCTCCGCCGGGCCAGCTCATTCTTAAAGTTCTAATACCGGGGTACGCTGCGGGGGCGGTGATCGGGAAAGGCGGTCAGATTATTGTACAACTTCAGAAAGATTCAGGGGCCATTATTAAGCTGTCAAAAGCGAAGGACTTTTACCCCGGAACCCAAGACCGAGTCGTTTTGATCCAAGGAACCGCCGAAGGCTTGATGAAGGTGCAAAATACCATTATAGAGAAGGTGTACGAGTTCCCTGTGCCCAAAGATTTAGCTGCGATCATCGGAGACCGACCGAAACAGGTGAAAATCATCGTACCCAACACAACTGCGGGACTGGTAATAGGAAAGGCCGGCGCAACGATAAAGACCATTATGGAAGAGAGTGGATCGAAGGTTCAACTCTCGCAAAAGCCAGACGGGGTAAACGTCCAAGAACGAGTCATCACAATCAAAGGAGAGAAGCACCAACTCATGACAGCATCTAATATTATTATTGATAAAATTAAAGACGACCCTCAAAGCGCCAGTTGCCCTCACATAAGTTACTCTGGCATCGCTGGCCCGATCGCTAACGCGAATCCCACCGGATCGCCCTACGCTGCTGGCTCGGCTGCATTAGTTGACGCTTCGCACCCATCCGTGGCCGCTATGTTGGGACATTATGTTATCCCAGGCCAACAGGTGCTGCAGACAGCAATGCCACTCTCCCATCACCCGCACCAGTCCGCGTTGTCCAGCGGCTCAGTGACACCGGCGCCTGAACTGACGACCATAAATCACGCCATGACAACGTTAGCGAACTATGGCTACACCTTAGGAGGCGTGAACTATGGTACCTTGGGTGTAATGCCTAGTGTACATCCAAGTGTACACCCTGGCATCGCTACCTCGGTCGGGATGATCTCTGCAGGCTCCCTAGCAGGAAGTCCAATCCCTTCAGCTACCCCCTTGCTCTCTGCCACTGCTCTACCGACGGAATCCAGTATTCCGACGGCTGTTCCCACTGCCCAAGCCATTTCAATGCAGAGCAATTACCTTGCAAACTTGGCTAATGCTGGTTACCTGACTACCGGTCACCCACAGTTGCTTGGAGCGACTATCCTAAGCATCGAAAAGTCAAGCGACGGACAAAAAGAAACAATTGAACTGGCAATTCCCGAAAACCTGATCGGAGCAGTCCTCGGAAAAGCGGGAAGGACACTGGTTGAGTATCAGGATGTATCAGGGGCGAAAATTCAAATTTCTAAAAAGGGTGATTACGTCGCCGGGACCAGGAACAGGAGGGTTACGATTACGGGGAAGCCCCCATGCCCACAGACTGCGCAGTTTCTTATTACGCAACGTGTCGCCTCTGCGCAAAACGCAAGGGCACAGCAGGCTAAGTTACTGTAGgaattc

*Islet -7216/-3950 + bpFOG>Nova(MLN) rescue*

Islet -7216/-3950 (Stolfi et al. 2010)

bpFOG (Rothbächer et al. 2007)

Nova(MLN) rescue

Silent mutation in Nova2.1 target sequence

ggcgcgccaagtataccacgcgagttagttcggccaggaagcgcgctgtttattatcaagttattatgacgtcatataaaatacgtcacatttgtagcacctggttatctgccttcccacggttaaaacagatgcgacctcggttatcagtggtttgacatggagatcgtaaatcaagcgacgaatatctgtgacacgtcacgtgacgagattgctgacataccaggacacttaacggtatctggttaggggaagtacctgtagcgtcatacacataacctgcataatgctttatagcttcgaatacgatagcgaagatcaaattaattaaaacaacgaagagaaggattcgaagcaaaacgacagtcgttaaaacacactacgggtaaaaattacttggggaaaagcgtggctgttacatccaaatcaaaatgtaagccgtcaatggtttgtgaacaaacggtaactcttgtcacctctaagctaaactcattttttgcagagtttaaaaataccgagtagcaactgccgtctggaacgaataaatgaatctaacgacgcagacgttatgacaggtgtgttcatacaccatgtttcccttttacgggaaagcgtataaaatatatatttacattcttgcctaaagtataaacgaacttttattgaatggtcaaaatgggtcatcttaaattccattataacttttccatttcaaacatttttccgcagcatcgtcataaaacagggattccaatagaaacatcataaaacaccaaaccatgttgtttatggatcaccgtgtttgttgtattaacaagagttataactacaacatgacgtgactagattgacacaacaactatggttggcagtcaaaacacacgaccccaccccataagatggagccacgccaccaaatcaacacagcagaagctccaggtggtttggtaggttctgttatcacaacactacaggccatgagatctatgatctataccgtctgttatatttagtggatttgtgattttaaccgaacaccttcactagtttaaaccaaccgacaactcaccgcgaacaatcagcgaatatttcattaaaccgacaaaaatttccagttaatttaaaatatagtaagttggggtaagatgggacatgtttttattctcatctcgtcccatttggtagtcaacaaagaatatttacagaattataaattgaaatatcttcacgaagagaaaaatattgttatttgttaaaaacacgatcaggaaatatgggatattttgtgctaaccgtattcatatttgcaacccactgtgagttgcttcttataaggtttaacagccacggttttaactaaattttatttattttaaccttagcgtgttataacgactgttatttttctgttaattcccatttcttcgccatgttagttaatatgacttttgttattgtaattatataaaaatatattttttaaaataagtctaaaagtaaatgtctaagttgtatttacaaatttccgcaccagaataggcagcaggcaaccgcggatcgaaacaagtccctgatttatggttttgttgcaactaattatggctgtacagcacacaggtcccaggattgccgtaaaacttaatttgctggaatatgttgaacaacagttgttacattctgttgttttattaagttcagcggccacgcttttttaagccttttattccagtattttcgctcgtagcgtgttgataacgactgttgttttgctgctaattctcttcgttgttttaattaaacttcgctatcgtattcaaaatatgttcatttagttggtaggttatgtgtgtaagtgacatcataacaacttgtatgaggcaaacaaggggttatgacgtaataatagaattattgatgtttcacagggtgaggtgtgacgtaataatagttctgacagttaaagtgattttgaataaaaacaaaaaacagaatgtttaagtcgcgaaaatcgcgtgggcagcgttaaccagagcatcgcatgtgacaagcgaacgaaatcccagcagatctacgacgttggtccgctgagagttcgagcgaattcgggatgtaacgtcgatgaaagacgaaaagcggtccttgcctcgacaagaaaggacgaaccgcgcgatagtagattttatggcgtatggtgttctataacacagctgtgggttgaagtggtttcgaatatttaagcaaggtacgacttgaccggttacgtcactttgaccacgtggtcagctctcatcaatatttattttaaatttgattaaacagcgacctcaggtgagcggcctaattatgcgcctctatgacgtcatagtttaaacttcttttaatagcggaatcgaagataaataggttgcgttaccctgggggtcgagatccaaatttcgttacttcgattgtttaattgattcgacatcgagttaaagcgcgagatttgattacatttccattgttcaatccataattcgctttgtgacgtaacacaaacttaacaaacacgtggtagcgcgcaagagctaattttttaaacgtttcttgattttttttcgattttttttattaaaaaataaaataaattaatattttgttgaaatacgtcgaaaccgcgaaaacgcgatcgacgcaattcaatttgtaaatgaaatatcgaccagtggcgccgatagcggataatcctcacaattcttgtgataaattagtttcttactcggataatctgtgtgccggtttgattgtcgcacccaaggtgattggggggggggggatattacgtcattatgacaaataatttcgattaaaattggtaaaaatataaattaggggagtttttagttttgttttaaaagattttatttaaaaattaaaaaaaaaatcgaaaatttttgaatttgatcgattagctgtaattacgtcatcaatccgctcattatgacgtcatacgtcgccatattgcgtcataatcgggcgtcagaatgtcgtcacaaaccatattgtgttgaattaattggaattatgaaacgccgctcggtcgggggatttgtgacgtcacaaacgtgaatgttacgacacaaacggcgaactcgggcgaacttttagctttgcggataataagggtaagtgacgcaacaatgacgcaacaatgttgtcccttgtgacgtaacagagtgtattgtaaccgttgaattgagtctcgagcagctgaagcttgcatgcctgcaggtcgactctagaggatccggcaaAGCTTCGTGTATTGTACCGGCCCATTGTCAATCATGCAAACTTGATATTATATTGACAAGAGAAGAAGGCAGTTTAAATTAAAACTCTAAAGTAGAGAGACATTAATCTCAGCTGACAAGGCAGGTGGTCACAGTAAGTTCATTTAAATAGTTGGCCAACAATAGCCTTTCCAAGAAAGTATTTTTGTTCCAGGTCTATACAAAAATAACACACATAgcggccgcaaccATGCTAAATGCAATGGAGTATGAATGCCAGTACAATGCTGGCTACAGCATTGTGTCTAACGGTAACGAATACGGTCTCATACAGGCCTACACGGCACACGATTACCCCCTTGAAAACGGAGTGACGTTTTCAGCACCTCCGCCcGGCCAGCTCATTCTTAAAGTTCTAATACCGGGGTACGCTGCGGGGGCGGTGATCGGGAAAGGCGGTCAGATTATTGTACAACTTCAGAAAGATTCAGGGGCCATTATTAAGCTGTCAAAAGCGAAGGACTTTTACCCCGGAACCCAAGACCGAGTCGTTTTGATCCAAGGAACCGCCGAAGGCTTGATGAAGGTGCAAAATACCATTATAGAGAAGGTGTACGAGTTCCCTGTGCCCAAAGATTTAGCTGCGATCATCGGAGACCGACCGAAACAGGTGAAAATCATCGTACCCAACACAACTGCGGGACTGGTAATAGGAAAGGCCGGCGCAACGATAAAGACCATTATGGAAGAGAGTGGATCGAAGGTTCAACTCTCGCAAAAGCCAGACGGGGTAAACGTCCAAGAACGAGTCATCACAATCAAAGGAGAGAAGCACCAACTCATGACAGCATCTAATATTATTATTGATAAAATTAAAGACGACCCTCAAAGCGCCAGTTGCCCTCACATAAGTTACTCTGGCATCGCTGGCCCGATCGCTAACGCGAATCCCACCGGATCGCCCTACGCTGCTGGCTCGGCTGCATTAGTTGACGCTTCGCACCCATCCGTGGCCGCTATGTTGGGACATTATGTTATCCCAGGCCAACAGGTGCTGCAGACAGCAATGCCACTCTCCCATCACCCGCACCAGTCCGCGTTGTCCAGCGGCTCAGTGACACCGGCGCCTGAACTGACGACCATAAATCACGCCATGACAACGTTAGCGAACTATGGCTACACCTTAGGAGGCGTGAACTATGGTACCTTGGGTGTAATGCCTAGTGTACATCCAAGTGTACACCCTGGCATCGCTACCTCGGTCGGGATGATCTCTGCAGGCTCCCTAGCAGGAAGTCCAATCCCTTCAGCTACCCCCTTGCTCTCTGCCACTGCTCTACCGACGGAATCCAGTATTCCGACGGCTGTTCCCACTGCCCAAGCCATTTCAATGCAGAGCAATTACCTTGCAAACTTGGCTAATGCTGGTTACCTGACTACCGGTCACCCACAGTTGCTTGGAGCGACGTCAGGCCTCGGCGGTCTCACCACAGTGTCCCAGCACCCGCCACCAGCGGCGACACCAACGAGTTTTTCCGTCGCTTCTACCCCTTCTACCCCTGGTCTGCCGGTTTCATTTAGCCCCCATTCAACCGTGAGTATCCTAAGCATCGAAAAGTCAAGCGACGGACAAAAAGAAACAATTGAACTGGCAATTCCCGAAAACCTGATCGGAGCAGTCCTCGGAAAAGCGGGAAGGACACTGGTTGAGTATCAGGATGTATCAGGGGCGAAAATTCAAATTTCTAAAAAGGGTGATTACGTCGCCGGGACCAGGAACAGGAGGGTTACGATTACGGGGAAGCCCCCATGCCCACAGACTGCGCAGTTTCTTATTACGCAACGTGTCGCCTCTGCGCAAAACGCAAGGGCACAGCAGGCTAAGTTACTGTAGgaattc

*Vacht -2083/+15>Unc-76::mCherry*

Vacht (Slc18a3, Cirobu.g00000742) -2083/+15 (Based on Yoshida et al. 2004)

Unc-76::mCherry

GGCGCGCCattacgtcgtaaacctttggctaccatcatctgcctcaaaacaaaattaattaaagaaatgcgttagtgtatcctttgactcggaatcaatcaaatcaagcaaaaatcaatatgtgaaattaaccatttagaccttgtgtcattcccattgcggtgacttgtcctagttgtgcgtttttatcagcagattgatttaacagtcaccggaacaggcaagacaatatttcaaaccagccaatgttaatttcagacaaatgaagcaatctgaaaatcagaacaaatcaaaaaacataagttttggtttttaaagcatagaaaacgtaccgtattttttaatgtaattgttaaattttgttatttaatatagtagggtagggggagatgggacactttttcattctgttttctcgtcttggtagcaaacaaaaacattaaaagaattataaaaccgtatcctcgcgactcctacagaccgtttaaaacaggatatttggatattctgttctaaaggtgtcccatcttaccccacagtactatacactatacacattctgtaccgatatattttattgattaaatttgaagttgttaaacttaattacgattaaatttcggcaaattgaaaatgagccatgaattaatcaaaaattattcttgatcgttgttttgtaaaacataacttttttttgatttttttgggaggcgccctgtgttccacttattattatttgtttgttttgctatgtcaatactttaatataaaagaaattaatattggctgacatttcaatttaacgctaggcttatgttttgtttcgtgaataatctgcataaagaaaaaaagcagtatcgactctcctattgttgaatcactcttgcttcccttccattgtgcaggagatagtgacgcaatgaacattgattttaacgcttctagtcagttgggcccttgctcacaattgcctaaaatttgatcaattgtggattgaaaagttaatattctttcctgaattaacatgtctataggtaaggtattgagacggctcccatttaatttcttgtccgatctttataaaaaaatatttgttaaaggtacgtttaatttgaacatatagtgcagaagtcttcggaattttcatatagagttattttttagatactatatgcgacatattttcatagcccaaattaaattgttttataatttgtttggcttttacagattttaatttaacttaggtgtacgattcaatagaaagattttaacctgtaaaaatacgtgcaagttgtttggaaataccttttttgttggacaaagttgtatcaggtaagttgatagcatgtatcttaaatctggcgtggtgtttatgtattggcttcatcatgaaatagtttgttttgtccttttacttgtcaattttatttaaattacatgagggtacaattcattataacgtacttcggtgaaaggttaatgttaacaaatgccgtccggtcttctgcgtatctttcgatttcgtattaattatgcatagaaagggtttaagtgcaatgctatttctgatgatgtgtgtgtaacaccaacgacctgatgacgaaaaacttgactgtttattaaataaaatagcgcaacaagacggcggaatttataaatggcatgttgctcgtaaacagcagcgcgtgcctgttttaactctggagaaacaagttatattcaactatttgttatctcacaaagcacggtattgcactcttatctaatgtactaatacaaatatatatatatatatatatatatatatctagctgttgtcgaggtaagatgaactgcagttaacaataaaaaccaacttcgttatacacaaattcaatataaagcgacagctcagtgttgtagagtaggtacccaactttcttaatctgcaaaaggcaaatacatgatttataatatgtaactcagcataacaacctgtttttgccttttgcagattacatctgagggcgggttgtgtgggtcagaaatttatcaaaggaaaataaagttgattgaaagcaaatttgtttctactttattgttcatcatgGACGTTTGTAGAgcggccgcaaccATGGCGGATCTGCGAGTACCGGACATTCCGCTCGCCTCGTGTGATGATGATGATATCGATAGTAATAAGAATTTGAGCAACCATTCATCAGACGAGAAACATCACTGCAACAGCAACAGCGACGAGGAACGTCTTCATGACGAGTTCTCTGGATCCCTTGAGGACCTTGTCGGCAACTTTGACGAAAAAATTGCGGCATGCCTGAAGGACCACGAGGTGACGACAGCGGATATTGCACCTGTGCAGATACGTACTCAAGAGGAAGTTATGAATGAAAGCCAAACATGGTGGACATTAACCGGAAACTTTGGAAACATTCAACCTCTCGACTTTGGAACCTCTTCGATATGTAAAAAGATGGCCGCAGCTCTGGACAGTGATTCATTGAAAGACGACGCATCTACACGCCGAAGTATGACAAATTCCGATGATGAGGATCTTTTACGACAACAAATGGATGTTCATCAAATGATTGGACATCATCATGGATCTACGGATACTGGTGGTGAAACACCTCCACAGACTGCTGATCAAGTTATCGAAGAAATTGATGAAATGTTACAGGTACCGGTCGCCACCATGGTGAGCAAGGGCGAGGAGGATAACATGGCCATCATCAAGGAGTTCATGCGCTTCAAGGTGCACATGGAGGGCTCCGTGAACGGCCACGAGTTCGAGATCGAGGGCGAGGGCGAGGGCCGCCCCTACGAGGGCACCCAGACCGCCAAGCTGAAGGTGACCAAGGGTGGCCCCCTGCCCTTCGCCTGGGACATCCTGTCCCCTCAGTTCATGTACGGCTCCAAGGCCTACGTGAAGCACCCCGCCGACATCCCCGACTACTTGAAGCTGTCCTTCCCCGAGGGCTTCAAGTGGGAGCGCGTGATGAACTTCGAGGACGGCGGCGTGGTGACCGTGACCCAGGACTCCTCCCTGCAGGACGGCGAGTTCATCTACAAGGTGAAGCTGCGCGGCACCAACTTCCCCTCCGACGGCCCCGTAATGCAGAAGAAGACCATGGGCTGGGAGGCCTCCTCCGAGCGGATGTACCCCGAGGACGGCGCCCTGAAGGGCGAGATCAAGCAGAGGCTGAAGCTGAAGGACGGCGGCCACTACGACGCTGAGGTCAAGACCACCTACAAGGCCAAGAAGCCCGTGCAGCTGCCCGGCGCCTACAACGTCAACATCAAGTTGGACATCACCTCCCACAACGAGGACTACACCATCGTGGAACAGTACGAACGCGCCGAGGGCCGCCACTCCACCGGCGGCATGGACGAGCTGTACAAGTAAgaattc

*Nova[MLN]-2011/+6>mScarlet*

Nova[MLN]-2012/+6

mScarlet (Bindels et al. 2017)

ggcgcgccTCAAAAGCCCATACTGCTCTATATAGCGCTCTTTTGGTTTATAAATTAGGAAAACTCTGTCGTGCCAGTTTAGTAAATTGCGATTTTTTGGTTCACTGCGGTTGTCTTACACGGTTAATGTAACTAACCGTGTAATGTATTTCACGAAGAATTAAAAACAAGCAACGTTTTGACTTCTGTGTAGAGTGTGCTCTTCATTCGAAGTGATGTGTAGGTAGTTTTGTATCGATTATATGCTCTGGCTGGGCCATATGAAAGCCCAGCTCGATCTGCATGTTTGTCAACAAGTTACATAGCACAAACAACGCTAGACGGATGCACTTGACCGAACTTTAATACTGATTAGATAAGAAAAAAACGGCAAAATGTGCAATACTACATTGGGATCGATGCGTTTCGATTGTGACAATGAGCAAAATTCCGATAATTAAAAGTAGTTGATATTTTTTTAATACAAAACTATCGAATTAATTACAAAAGGTACCACATTCTTATACCAAAGCATTTAGATTTATTGATTTGAAGAATGTATGCGATCGAAACGAAGCAATTTCCTGATATAGTTAAAAGGTCATCAGAATAAATGGAAACTTAGTGACGTCGACCCCAATCATACGCTCGCAATTTTGGTCCATCGCTAATGCGCCTTTGGGAAATCATGGGACGACATCACGAGAGATTAGTTTCGAACTGCTTTGGTCTCTGCAGTGTTTCCAGACGGTTTCTATTTCGCACCCGGAAACACTGATTCGCCTTAGCGCGGCTTAGTGGGTTCGTCATTAAGAAATCGAAGTGAGATTGTTTACCAAACACATGCGCTTAGTTGTCTGTGGTTTCTGCCGTCGTATCATTAATTTGTATACATTTGGTAACAAATGCGAAATGACAGCGATTCGGACCGATTTTTTTCATCAAATTTTAACTGCGTGTTTCACATTATTTCTTAAAGCACGTCCGTGTTGCAATGCTTTGTAATGGATTATATTTGTGAGGCTGTAAATTACAAACCAGCGGCGGTACTTCAATTCCAGCACATGGTCGTAAAGAATTGTCCGAACATTACTGCGGGTAGCTCTGTACGTTTCGCCAATGATACTCTCAGTTAATGTTCAGCCATTAGTCGGGCATGCGATGCCAAAGTTTGCTCGGCAGTAAAATTGGATGCTTTGAAAAAAATCTAATAATTAACTGAAGCGGTTGTAGAACCGTCGTAAAAGTACCGCACTGAATGTAAAGGGGTGCAGAGGCTGACTACGCTAGACAAAGGTGCTGGTTTTACGACAAGAGTGCTTCCCGATTGCGATACGAAAAAAACAGTATTGTTGCGAGAGGGTGCGGTTAAAAAATGATTTTGAAACACTGTTTTCCGGAGTGCTGAGCTGTTCAAACGCAGCGTACAGCAACAATGCTTGTGCAGTGATCGAGTTGCTACGCGTTCTGTTTGATGGCTTCTATAAATAAAATGAAAGTTGTTGGTACCAGTCCGCCCAGAAGCGGTGCTTATTCGTTCGTAATAGGCGGTGCATTTCAGGAAGCAATACGTCGAGTGTTTTTCTTGCTTTTCTCCCAGAAATCTAAGAACAATAAATTGAATAAAGGACGGCGACTGCTAGGGCAAGATATCTGGCCAGATTCGAAAGCGCAGGTCCGCATCTAGGTTGCATCGTTGCCGGAAAGAGTGGGCTGAGTCTATAATGTGTATCCACGCGCGCAGTGATGTTGGAAGGCCCTGTTAGTATCAGCTCATTGTATGCAGCAGTGTTCTCGCCTGATTGACTGCGTTTGAGGTTAAAATTTAATGGCGTTGCGGTTTTCCTGTTTCTATGAACTGAGGTCGCAAGCAATTATAGATGTATTCGATCGATGAATATGAAAAGGGGCCGTATATTGTTGAGTGATACGTGTAAGGGTTAAAGCTTGTAGTTATATAAGTACAGAAGAGCCTGACGTATTGGAGGTTCATGATTTAATCATTAATCGCTCTTTTCATTCTGACAGAGAAAGTAGGATAATGCTAgcggccgcaaccATGGCTAGCGTGAGCAAGGGCGAGGCAGTGATCAAGGAGTTCATGCGGTTCAAGGTGCACATGGAGGGCTCCATGAACGGCCACGAGTTCGAGATCGAGGGCGAGGGCGAGGGCCGCCCCTACGAGGGCACCCAGACCGCCAAGCTGAAGGTGACCAAGGGTGGCCCCCTGCCCTTCTCCTGGGACATCCTGTCCCCTCAGTTCATGTACGGCTCCAGGGCCTTCACCAAGCACCCCGCCGACATCCCCGACTACTATAAGCAGTCCTTCCCCGAGGGCTTCAAGTGGGAGCGCGTGATGAACTTCGAGGACGGCGGCGCCGTGACCGTGACCCAGGACACCTCCCTGGAGGACGGCACCCTGATCTACAAGGTGAAGCTCCGCGGCACCAACTTCCCTCCTGACGGCCCCGTAATGCAGAAGAAGACAATGGGCTGGGAAGCGTCCACCGAGCGGTTGTACCCCGAGGACGGCGTGCTGAAGGGCGACATTAAGATGGCCCTGCGCCTGAAGGACGGCGGTCGCTACCTGGCGGACTTCAAGACCACCTACAAGGCCAAGAAGCCCGTGCAGATGCCCGGCGCCTACAACGTCGACCGCAAGTTGGACATCACCTCCCACAACGAGGACTACACCGTGGTGGAACAGTACGAACGCTCCGAGGGCCGCCACTCCACCGGCGGCATGGACGAGCTGTACAAGTAAgaattc

*Nova[MLN]-2011/+6* driver (wild-type)

tcaaaagcccatactgctctatatagcgctcttttggtttataaattaggaaaactctgtcgtgccagtttagtaaattgcgattttttggttcactgcggttgtcttacacggttaatgtaactaaccgtgtaatgtatttcacgaagaattaaaaacaagcaacgttttgacttctgtgtagagtgtgctcttcattcgaagtgatgtgtaggtagttttgtatcgattatatgctctggctgggccatatgaaagcccagctcgatctgcatgtttgtcaacaagttacatagcacaaacaacgctagacggatgcacttgaccgaactttaatactgattagataagaaaaaaacggcaaaatgtgcaatactacattgggatcgatgcgtttcgattgtgacaatgagcaaaattccgataattaaaagtagttgatatttttttaatacaaaactatcgaattaattacaaaaggtaccacattcttataccaaagcatttagatttattgatttgaagaatgtatgcgatcgaaacgaagcaatttcctgatatagttaaaaggtcatcagaataaatggaaacttagtgacgtcgaccccaatcatacgctcgcaattttggtccatcgctaatgcgcctttgggaaatcatgggacgacatcacgagagattagtttcgaactgctttggtctctgcagtgtttccagacggtttctatttcgcacccggaaacactgattcgccttagcgcggcttagtgggttcgtcattaagaaatcgaagtgagattgtttaccaaacacatgcgcttagttgtctgtggtttctgccgtcgtatcattaatttgtatacatttggtaacaaatgcgaaatgacagcgattcggaccgatttttttcatcaaattttaactgcgtgtttcacattatttcttaaagcacgtccgtgttgcaatgctttgtaatggattatatttgtgaggctgtaaattacaaaccagcggcggtacttcaattccagcacatggtcgtaaagaattgtccgaacattactgcgggtagctctgtacgtttcgccaatgatactctcagttaatgttcagccattagtcgggcatgcgatgccaaagtttgctcggcagtaaaattggatgctttgaaaaaaatctaataattaactgaagcggttgtagaaccgtcgtaaaagtaccgcactgaatgtaaaggggtgcagaggctgactacgctagacaaaggtgctggttttacgacaagagtgcttcccgattgcgatacgaaaaaaacagtattgttgcgagagggtgcggttaaaaaatgattttgaaacactgttttccggagtgctgagctgttcaaacgcagcgtacagcaacaatgcttgtgcagtgatcgagttgctacgcgttctgtttgatggcttctataaataaaatgaaagttgttggtaccagtccgcccagaagcggtgcttattcgttcgtaataggcggtgcatttcaggaagcaatacgtcgagtgtttttcttgcttttctcccagaaatctaagaacaataaattgaataaaggacggcgactgctagggcaagatatctggccagattcgaaagcgcaggtccgcatctaggttgcatcgttgccggaaagagtgggctgagtctataatgtgtatccacgcgcgcagtgatgttggaaggccctgttagtatcagctcattgtatgcagcagtgttctcgcctgattgactgcgtttgaggttaaaatttaatggcgttgcggttttcctgtttctatgaactgaggtcgcaagcaattatagatgtattcgatcgatgaatatgaaaaggggccgtatattgttgagtgatacgtgtaagggttaaagcttgtagttatataagtacagaagagcctgacgtattggaggttcatgatttaatcattaatcgctcttttcattctgacagagaaagtaggataatgcta

*Nova[MLN]-2011/+6 mEBF 1*

tcaaaagcccatactgctctatatagcgctcttttggtttataaattaggaaaactctgtcgtgccagtttagtaaattgcgattttttggttcactgcggttgtcttacacggttaatgtaactaaccgtgtaatgtatttcacgaagaattaaaaacaagcaacgttttgacttctgtgtagagtgtgctcttcattcgaagtgatgtgtaggtagttttgtatcgattatatgctctggctgggccatatgaaagcccagctcgatctgcatgtttgtcaacaagttacatagcacaaacaacgctagacggatgcacttgaccgaactttaatactgattagataagaaaaaaacggcaaaatgtgcaatactacattgggatcgatgcgtttcgattgtgacaatgagcaaaattccgataattaaaagtagttgatatttttttaatacaaaactatcgaattaattacaaaaggtaccacattcttataccaaagcatttagatttattgatttgaagaatgtatgcgatcgaaacgaagcaatttcctgatatagttaaaaggtcatcagaataaatggaaacttagtgacgtcgaccccaatcatacgctcgcaattttggtccatcgctaatgcgcctttAAgaaatcatgggacgacatcacgagagattagtttcgaactgctttggtctctgcagtgtttccagacggtttctatttcgcacccggaaacactgattcgccttagcgcggcttagtgggttcgtcattaagaaatcgaagtgagattgtttaccaaacacatgcgcttagttgtctgtggtttctgccgtcgtatcattaatttgtatacatttggtaacaaatgcgaaatgacagcgattcggaccgatttttttcatcaaattttaactgcgtgtttcacattatttcttaaagcacgtccgtgttgcaatgctttgtaatggattatatttgtgaggctgtaaattacaaaccagcggcggtacttcaattccagcacatggtcgtaaagaattgtccgaacattactgcgggtagctctgtacgtttcgccaatgatactctcagttaatgttcagccattagtcgggcatgcgatgccaaagtttgctcggcagtaaaattggatgctttgaaaaaaatctaataattaactgaagcggttgtagaaccgtcgtaaaagtaccgcactgaatgtaaaggggtgcagaggctgactacgctagacaaaggtgctggttttacgacaagagtgcttcccgattgcgatacgaaaaaaacagtattgttgcgagagggtgcggttaaaaaatgattttgaaacactgttttccggagtgctgagctgttcaaacgcagcgtacagcaacaatgcttgtgcagtgatcgagttgctacgcgttctgtttgatggcttctataaataaaatgaaagttgttggtaccagtccgcccagaagcggtgcttattcgttcgtaataggcggtgcatttcaggaagcaatacgtcgagtgtttttcttgcttttctcccagaaatctaagaacaataaattgaataaaggacggcgactgctagggcaagatatctggccagattcgaaagcgcaggtccgcatctaggttgcatcgttgccggaaagagtgggctgagtctataatgtgtatccacgcgcgcagtgatgttggaaggccctgttagtatcagctcattgtatgcagcagtgttctcgcctgattgactgcgtttgaggttaaaatttaatggcgttgcggttttcctgtttctatgaactgaggtcgcaagcaattatagatgtattcgatcgatgaatatgaaaaggggccgtatattgttgagtgatacgtgtaagggttaaagcttgtagttatataagtacagaagagcctgacgtattggaggttcatgatttaatcattaatcgctcttttcattctgacagagaaagtaggataatgcta

*Nova[MLN]-2011/+6 mEBF 2*

tcaaaagcccatactgctctatatagcgctcttttggtttataaattaggaaaactctgtcgtgccagtttagtaaattgcgattttttggttcactgcggttgtcttacacggttaatgtaactaaccgtgtaatgtatttcacgaagaattaaaaacaagcaacgttttgacttctgtgtagagtgtgctcttcattcgaagtgatgtgtaggtagttttgtatcgattatatgctctggctgggccatatgaaagcccagctcgatctgcatgtttgtcaacaagttacatagcacaaacaacgctagacggatgcacttgaccgaactttaatactgattagataagaaaaaaacggcaaaatgtgcaatactacattgggatcgatgcgtttcgattgtgacaatgagcaaaattccgataattaaaagtagttgatatttttttaatacaaaactatcgaattaattacaaaaggtaccacattcttataccaaagcatttagatttattgatttgaagaatgtatgcgatcgaaacgaagcaatttcctgatatagttaaaaggtcatcagaataaatggaaacttagtgacgtcgaccccaatcatacgctcgcaattttggtccatcgctaatgcgcctttgggaaatcatAAgacgacatcacgagagattagtttcgaactgctttggtctctgcagtgtttccagacggtttctatttcgcacccggaaacactgattcgccttagcgcggcttagtgggttcgtcattaagaaatcgaagtgagattgtttaccaaacacatgcgcttagttgtctgtggtttctgccgtcgtatcattaatttgtatacatttggtaacaaatgcgaaatgacagcgattcggaccgatttttttcatcaaattttaactgcgtgtttcacattatttcttaaagcacgtccgtgttgcaatgctttgtaatggattatatttgtgaggctgtaaattacaaaccagcggcggtacttcaattccagcacatggtcgtaaagaattgtccgaacattactgcgggtagctctgtacgtttcgccaatgatactctcagttaatgttcagccattagtcgggcatgcgatgccaaagtttgctcggcagtaaaattggatgctttgaaaaaaatctaataattaactgaagcggttgtagaaccgtcgtaaaagtaccgcactgaatgtaaaggggtgcagaggctgactacgctagacaaaggtgctggttttacgacaagagtgcttcccgattgcgatacgaaaaaaacagtattgttgcgagagggtgcggttaaaaaatgattttgaaacactgttttccggagtgctgagctgttcaaacgcagcgtacagcaacaatgcttgtgcagtgatcgagttgctacgcgttctgtttgatggcttctataaataaaatgaaagttgttggtaccagtccgcccagaagcggtgcttattcgttcgtaataggcggtgcatttcaggaagcaatacgtcgagtgtttttcttgcttttctcccagaaatctaagaacaataaattgaataaaggacggcgactgctagggcaagatatctggccagattcgaaagcgcaggtccgcatctaggttgcatcgttgccggaaagagtgggctgagtctataatgtgtatccacgcgcgcagtgatgttggaaggccctgttagtatcagctcattgtatgcagcagtgttctcgcctgattgactgcgtttgaggttaaaatttaatggcgttgcggttttcctgtttctatgaactgaggtcgcaagcaattatagatgtattcgatcgatgaatatgaaaaggggccgtatattgttgagtgatacgtgtaagggttaaagcttgtagttatataagtacagaagagcctgacgtattggaggttcatgatttaatcattaatcgctcttttcattctgacagagaaagtaggataatgcta

*Nova[MLN]-2011/+6 mEBF 1+2*

tcaaaagcccatactgctctatatagcgctcttttggtttataaattaggaaaactctgtcgtgccagtttagtaaattgcgattttttggttcactgcggttgtcttacacggttaatgtaactaaccgtgtaatgtatttcacgaagaattaaaaacaagcaacgttttgacttctgtgtagagtgtgctcttcattcgaagtgatgtgtaggtagttttgtatcgattatatgctctggctgggccatatgaaagcccagctcgatctgcatgtttgtcaacaagttacatagcacaaacaacgctagacggatgcacttgaccgaactttaatactgattagataagaaaaaaacggcaaaatgtgcaatactacattgggatcgatgcgtttcgattgtgacaatgagcaaaattccgataattaaaagtagttgatatttttttaatacaaaactatcgaattaattacaaaaggtaccacattcttataccaaagcatttagatttattgatttgaagaatgtatgcgatcgaaacgaagcaatttcctgatatagttaaaaggtcatcagaataaatggaaacttagtgacgtcgaccccaatcatacgctcgcaattttggtccatcgctaatgcgcctttAAgaaatcatAAgacgacatcacgagagattagtttcgaactgctttggtctctgcagtgtttccagacggtttctatttcgcacccggaaacactgattcgccttagcgcggcttagtgggttcgtcattaagaaatcgaagtgagattgtttaccaaacacatgcgcttagttgtctgtggtttctgccgtcgtatcattaatttgtatacatttggtaacaaatgcgaaatgacagcgattcggaccgatttttttcatcaaattttaactgcgtgtttcacattatttcttaaagcacgtccgtgttgcaatgctttgtaatggattatatttgtgaggctgtaaattacaaaccagcggcggtacttcaattccagcacatggtcgtaaagaattgtccgaacattactgcgggtagctctgtacgtttcgccaatgatactctcagttaatgttcagccattagtcgggcatgcgatgccaaagtttgctcggcagtaaaattggatgctttgaaaaaaatctaataattaactgaagcggttgtagaaccgtcgtaaaagtaccgcactgaatgtaaaggggtgcagaggctgactacgctagacaaaggtgctggttttacgacaagagtgcttcccgattgcgatacgaaaaaaacagtattgttgcgagagggtgcggttaaaaaatgattttgaaacactgttttccggagtgctgagctgttcaaacgcagcgtacagcaacaatgcttgtgcagtgatcgagttgctacgcgttctgtttgatggcttctataaataaaatgaaagttgttggtaccagtccgcccagaagcggtgcttattcgttcgtaataggcggtgcatttcaggaagcaatacgtcgagtgtttttcttgcttttctcccagaaatctaagaacaataaattgaataaaggacggcgactgctagggcaagatatctggccagattcgaaagcgcaggtccgcatctaggttgcatcgttgccggaaagagtgggctgagtctataatgtgtatccacgcgcgcagtgatgttggaaggccctgttagtatcagctcattgtatgcagcagtgttctcgcctgattgactgcgtttgaggttaaaatttaatggcgttgcggttttcctgtttctatgaactgaggtcgcaagcaattatagatgtattcgatcgatgaatatgaaaaggggccgtatattgttgagtgatacgtgtaagggttaaagcttgtagttatataagtacagaagagcctgacgtattggaggttcatgatttaatcattaatcgctcttttcattctgacagagaaagtaggataatgcta

**Electroporation mixes (per 700 µl of solution):**

Testing sgRNA combinations for Agrin CRISPR by RT-PCR (Fig5B)

40 µg Eef1a>Cas9

60 µl bead-purified U6>Agrin sgRNA OSO-PCR products (10 µl each sgRNA)

Control sgRNA to compare to Agrin CRISPR by RT-PCR (Fig5B)

40 µg Eef1a>Cas9

50 µl bead-purified U6>Control sgRNA OSO-PCR product

Neural-specific Z+ Agrin CRISPR to see AChRA1::GFP clustering (Fig 5C)

40 µg Sox1/2/3>Cas9

60 µl bead-purified U6>Agrin sgRNA OSO-PCR products (10 µl each sgRNA)

40 µg Vacht -2083/+15>Unc-76::mCherry

20 µg Tbx6-r.b>AChRA1::GFP

Neural-specific control CRISPR to compare to Z+ Agrin CRISPR (Fig5C)

40 µg Sox1/2/3>Cas9

50 µl bead-purified U6>Control sgRNA OSO-PCR product

40 µg Vacht -2083/+15>Unc-76::mCherry

20 µg Tbx6-r.b>AChRA1::GFP

Neural-specific control CRISPR to compare to Agrin CRISPR 1+3 (Fig5D)

40 µg Sox1/2/3>Cas9

50 µg U6>Control

40 µg Vacht -4315/+15>Unc-76::mCherry

20 µg Tbx6-r.b>AChRA1::GFP

Neural-specific Z+ Agrin CRISPR sgRNAs 1+3 (Fig5D)

40 µg Sox1/2/3>Cas9

25 µg U6>Agrin.1

25 µg U6>Agrin.3

40 µg Vacht -4315/+15>Unc-76::mCherry

20 µg Tbx6-r.b>AChRA1::GFP

Neural-specific control CRISPR to compare to Agrin CRISPR 5+8 (Fig3D)

40 µg Sox1/2/3>Cas9

50 µg U6>Control

40 µg Vacht -4315/+15>Unc-76::mCherry

20 µg Tbx6-r.b>AChRA1::GFP

Neural-specific Z+ Agrin CRISPR sgRNAs 5+8 (Fig3D)

40 µg Sox1/2/3>Cas9

25 µg U6>Agrin.5

25 µg U6>Agrin.8

40 µg Vacht -4315/+15>Unc-76::mCherry

20 µg Tbx6-r.b>AChRA1::GFP

Muscle-specific Lrp4 CRISPR (Fig3E,F)

35 µg Tbx6-r.b>Cas9

60 µg Tbx6-r.b>CD4::mCherry

25 µg U6>Lrp4.2

25 µg U6>Lrp4.4

20 µg Tbx6-r.b>AChRA1::GFP

Muscle-specific Nova (negative control) CRISPR (Fig3E,F)

35 µg Tbx6-r.b>Cas9

60 µg Tbx6-r.b>CD4::mCherry

50 µg U6>Nova2.3

20 µg Tbx6-r.b>AChRA1::GFP

Nova CRISPR for RT-PCR, sgRNA 1.2(Fig6B)

40 µg Sox1/2/3>Cas9

40 µg U6>Nova1.2

Nova CRISPR for RT-PCR, sgRNA 2.1(Fig6B)

40 µg Sox1/2/3>Cas9

40 µg U6>Nova2.1

Nova CRISPR for RT-PCR, sgRNA 2.3(Fig6B)

40 µg Sox1/2/3>Cas9

40 µg U6>Nova2.3

Control CRISPR for RT-PCR, sgRNA 1.2(Fig6B)

40 µg Sox1/2/3>Cas9

40 µg U6>Control

Nova sgRNA “Mix” CRISPR for RT-PCR (Fig6B,C)

25 µg Ef1a>Cas9

25 µg U6>Nova1.2

25 µg U6>Nova2.1

25 µg U6>Nova2.3

Control CRISPR for RT-PCR (Fig6C)

25 µg Ef1a>Cas9

75 µg U6>Control

Eef1a>Nova(MLN) for RT-PCR (Fig6C)

20 µg Eef1a>Nova(MLN)

Eef1a>H2B::CFP as control for Nova(MLN) overexpression RT-PCR (Fig6C)

20 µg Eef1a>H2B:CFP

Neural-specific Nova CRISPR to compare to rescue (Fig6E)

40 µg Sox1/2/3>Cas9

25 µg U6>Nova1.2

25 µg U6>Nova2.1

35 µg Islet -7216/-3950 + bpFOG>Unc-76::mCherry

20 µg Tbx6-r.b>AChRA1::GFP

Neural-specific control CRISPR to compare to rescue (Fig4E)

40 µg Sox1/2/3>Cas9

50 µg U6>Control

35 µg Islet -7216/-3950 + bpFOG>Unc-76::mCherry

20 µg Tbx6-r.b>AChRA1::GFP

Rescue of neural-specific Nova CRISPR (Fig4E)

40 µg Sox1/2/3>Cas9

25 µg U6>Nova1.2

25 µg U6>Nova2.1

35 µg Islet -7216/-3950 + bpFOG>Unc-76::mCherry

20 µg Tbx6-r.b>AChRA1::GFP

40 µg Islet -7216/-3950 + bpFOG>Nova(MLN) rescue

Neural-specific Nova CRISPR for cluster density (Fig4D,F)

40 µg Sox1/2/3>Cas9

15 µg U6>Nova1.2

15 µg U6>Nova2.1

15 µg U6>Nova2.3

40 µg Vacht -2083/+15>Unc-76::mCherry

20 µg Tbx6-r.b>AChRA1::GFP

Neural-specific control CRISPR for cluster density (Fig4D,F)

40 µg Sox1/2/3>Cas9

45 µg U6>Control

40 µg Vacht -2083/+15>Unc-76::mCherry

20 µg Tbx6-r.b>AChRA1::GFP

Neural-specific Ebf CRISPR to assay Nova>GFP expression (Fig7C,D)

35 µg Sox1/2/3>Cas9

40 µg U6>Ebf.C

60 µg Nova[MLN] -2011/+6>GFP

45 µg Islet -7216/-3950 + bpFOG>tagRFP

Neural-specific Control CRISPR to assay Nova>GFP expression (Fig7C,D)

35 µg Sox1/2/3>Cas9

40 µg U6>DenhT2

60 µg Nova[MLN] -2011/+6>GFP

45 µg Islet -7216/-3950 + bpFOG>tagRFP

WT vs WT (Fig7F-H)

60 µg Nova[MLN] -2011/+6>GFP

60 µg Nova[MLN] -2011/+6>mScarlet

WT vs mEbf1 (Fig7F,G)

60 µg Nova[MLN] -2011/+6 mEbf1>GFP

60 µg Nova[MLN] -2011/+6>mScarlet

WT vs mEbf2 (Fig7F,G)

60 µg Nova[MLN] -2011/+6 mEbf2 >GFP

60 µg Nova[MLN] -2011/+6>mScarlet

WT vs mEbf1+2 (Fig7G)

60 µg Nova[MLN] -2011/+6 mEbf1+2>GFP

60 µg Nova[MLN] -2011/+6>mScarlet
